# Mapping Typical and Altered Neurodevelopment with Sleep Macro- and Micro-Architecture

**DOI:** 10.1101/2022.12.15.520643

**Authors:** N Kozhemiako, AW Buckley, RD Chervin, S Redline, SM Purcell

**Affiliations:** Brigham and Women’s Hospital & Harvard Medical School, Boston, MA, USA; National Institute of Mental Health, NIH, Bethesda, MD, USA; Department of Neurology, University of Michigan, Ann Arbor, MI, USA; Beth Israel Deaconess Medical Center, Boston, MA, USA

## Abstract

Profiles of sleep duration and timing and corresponding electroencephalographic activity reflect brain changes that support cognitive and behavioral maturation and may provide practical markers for tracking typical and atypical neurodevelopment. To build and evaluate a sleep-based, quantitative metric of brain maturation, we used whole-night polysomnography data, initially from two large National Sleep Research Resource samples, spanning childhood and adolescence (total *N* = 4,013, aged 2.5 to 17.5 years): the Childhood Adenotonsillectomy Trial (CHAT), a research study of children with snoring without neurodevelopmental delay, and NCH, a pediatric sleep clinic cohort. Among children without developmental disorders, sleep metrics derived from the electroencephalogram (EEG) displayed robust age-related changes consistently across datasets. Prominent stage-, band- and channel-specific developmental trajectories in spectral power were found. During non-rapid eye movement (NR) sleep, spindles and slow oscillations further exhibited characteristic developmental patterns, with respect to their rate of occurrence, temporal coupling and morphology. Based on these metrics in NCH, we constructed a model to predict an individual’s chronological age. The model performed with high accuracy (*r* = 0.95 in the held-out NCH testing sample and *r* = 0.88 in a second independent replication sample (PATS) with a broadly comparable age range). EEG-based age predictions reflected clinically meaningful neurodevelopmental differences; for example, compared to typically developing children, those with neurodevelopmental diagnoses (NDD) showed greater variability in predicted age, and children with Down syndrome or intellectual disability had significantly younger brain age predictions (respectively, 2.2 and 0.59 years less than their chronological age) compared to age-matched non-NDD children. Overall, our results indicate that sleep architecture offers a sensitive window for characterizing brain maturation, suggesting the potential for scalable, objective sleep-based biomarkers to measure typical and atypical neurodevelopment.

## Introduction

Brain structure and function undergo drastic transformations over the first two decades of life (Stiles & Jernigan, 2010). Over the same period, there are also profound changes in multiple facets of sleep, potentially reflecting developmental programs of brain reorganization (Knoop et al., 2021). Given abundant evidence for the critical impact that sleep has on memory consolidation and working memory, deviations from normal sleep patterns during childhood and adolescence – probably the most learning-intensive periods in life - might have long lasting consequences (Kopasz et al., 2010). For example, active processes taking place during sleep support synaptic pruning and connectivity restructuring (Buchmann et al., 2011; Tononi & Cirelli, 2014). A more precise characterization of normative childhood trajectories of sleep may therefore aid our understanding of brain developmental processes and shed light on how subsequent cognitive and behavioral dysfunction can emerge.

Children with neurodevelopmental disorders (NDD) and related diagnoses including epilepsy are exceptionally vulnerable to sleep problems (Kamara & Beauchaine, 2020; Robinson-Shelton & Malow, 2015). Sleep dysregulation has been documented in individuals with autism spectrum disorder (ASD) (Souders et al., 2017), attention deficit / hyperactivity disorder (ADHD) (Becker, 2020), cerebral palsy (Simard-Tremblay et al., 2011), Down syndrome (Stores & Stores, 2013), and epilepsy (Larson et al., 2012). Comorbid conditions that commonly accompany clinical manifestation of NDDs, including intellectual disability, have also been associated with sleep abnormalities (Surtees et al., 2018).

However, the majority of these reports focused on a single disorder despite evidence of trans-diagnostically shared risk factors and pathogenic mechanisms, especially for psychiatric disorders (Gandal et al., 2016; Thapar et al., 2017). A second limitation is that the most-studied aspects of sleep have been macro-level metrics (e.g. time in bed) approximated by parental report. Such measures are inherently limited with respect to characterizing brain activity. In contrast, electrophysiological characterizations of sleep micro-architecture reflect underlying processes during different sleep stages more directly. One recent review described the presence of sleep oscillation (e.g. spindles, slow oscillations) abnormalities among NDDs but also highlighted the scarcity of relevant studies and limited sample sizes which, combined with varying age ranges and inconsistent methodologies, precluded strong conclusions (Gorgoni et al., 2020).

Across the lifespan, when investigating age-related change numerous studies have adopted the paradigm of predicting chronological age based on neuroimaging or other data sources, on the assumption that its deviation from observed age – the *brain age gap* – is a putative marker of overall brain health. Compared to healthy controls, “brain age gaps” have been observed in various adult-onset disorders and conditions including Alzheimer disease, mild cognitive impairment, schizophrenia and multiple sclerosis (Baecker et al., 2021). Accelerated aging – larger positive brain age gaps – has also been shown to follow traumatic brain injury (Cole et al., 2015). Conversely, opposite patterns in children born very prematurely have been interpreted as reflections of delayed brain development (Franke et al., 2012). Although the majority of studies on brain age used structural MRI data, with predictions usually achieving high correlations (*r* > 0.9) with chronological age (Franke & Gaser, 2019), recent reports in adults demonstrated that the sleep EEG can also give comparable results (Nygate et al., 2021; Sun et al., 2019). In aggregate, these findings suggest that sleep EEG features track strongly with age and may be valuable for mapping typical and atypical childhood development.

In the present study, we used polysomnography (PSG) data from a large clinical cohort to comprehensively chart the developmental trajectories of metrics derived from sleep electroencephalograms (EEG) across childhood and adolescence. Based on clinical records, we initially removed individuals with any of six major clinical groups including NDDs and used data from remaining individuals (*N*=1,828, 2.5 – 17.5 years) to characterize broadly typical neurodevelopment. We further validated findings in an independent sample of generally healthy children with snoring. This latter group of children participated in a clinical trial and were screened to be free of severe neurodevelopmental delay (estimated by the Differential Ability Scales II), although they had a more restricted age range (*N*=1,213, 4.5 – 10 years). Based on the multivariate profile of observed associations, we developed a single model to predict chronological age as a function of sleep macro- and micro-architecture. Specifically, we tested 1) whether such a model was transferable between different studies and populations, and 2) whether deviations between predicted and chronological age distinguished children with NDD from typically developing children. For the last two goals we employed two additional datasets, from the Pediatric Adenotonsillectomy Trial for Snoring, (PATS, *N* = 627 children with snoring), and the Cleveland Family Study (CFS, *N* = 730), a family-based study covering a wide age range from 6 to 88 years.

## Results

The primary discovery dataset comprised 2,800 individuals with whole-night PSGs from the Nationwide Children’s Hospital (NCH) Sleep Databank (accessed via the National Sleep Research Resource, NSRR). Based on ICD codes, we defined six subsets within the NCH sample based on the presence of a diagnosis of the following neurodevelopmental and childhood onset disorders: ASD, ADHD, intellectual disabilities, Down syndrome (DS), cerebral palsy (CP) and epilepsy (for briefness below in the text, we use “NDD group” to refer to those sub-cohorts although we acknowledge that epilepsy is usually not classified as an NDD). We selected these six subgroups based on each having *N* > 100 subjects in the full dataset and previous reports of alterations in sleep (Becker, 2020; Larson et al., 2012; Simard-Tremblay et al., 2011; Souders et al., 2017; Stores & Stores, 2013; Surtees et al., 2018) (see **Table 2** for demographic details, and **Sup. Table 1** for diagnostic details). There was a considerable overlap between NDD diagnoses (**Sup. Figure 1a)**.

As expected in this clinically referred and ascertained sample, most NCH individuals had a sleep-related clinical diagnosis, precluding a straightforward definition of a “healthy control” comparison group. Although we will refer to the sample without NDD-subgroups as the non-NDD sample, individuals therein collectively had more than 9,000 unique diagnostic codes identified in their medical records, indicative of both acute and chronic disorders (although not necessarily contemporaneous with the PSG recording). To give a few examples, there are diagnoses of cough (*N*=1176), obesity (N=793), skin rash (*N*=633), unspecified disturbance of conduct (*N*=215) and anxiety disorder (*N*=210). Excluding the abovementioned six NDD groups, 93% of participants had one or more sleep-related diagnostic codes including sleep apnea, insomnia, hypersomnia and others. Sleep disorders were more prevalent among the clinical NDD sub-groups (**Sup. Figure 1b**). To control for medication effects, we further identified individuals across the NCH sample who were prescribed medications likely to affect sleep at a time overlapping the PSG recording. Only 14% (262 out of 1,829 non-NDD NCH individuals) of the NCH sample had such medication prescribed at the night of PSG, with a majority being antihistamines (9%) (**Sup. Figure 1c, Sup. Table 2**). The proportion of individuals prescribed sleep-impacting medication during PSG was substantially higher among the NDD subgroups (42%). Medication use was added as a covariate in our analyses (see Methods for more details).

For our initial analyses of typical age-related changes in sleep, we excluded the NDD groups resulting in a final sample of *N*=1,828 (detailed demographic information in **Table 1)**. For an independent replication cohort, we obtained PSGs from *N* = 1,213 individuals from the Child Adenotonsillectomy Trial (CHAT) study: children with reported snoring, screened to be without neurodevelopmental delay. The CHAT study had a narrower age range of 4.5 – 10 years compared to the NCH. We included children without obstructive sleep apnea OSA (Apnea Hypopnea Index, AHI<2), with mild-moderate OSA (2-30) as well as those with severe OSA (AHI >30) but excluded follow-up recordings after the adenotonsillectomy procedure (Table 1).

**Table 1.**
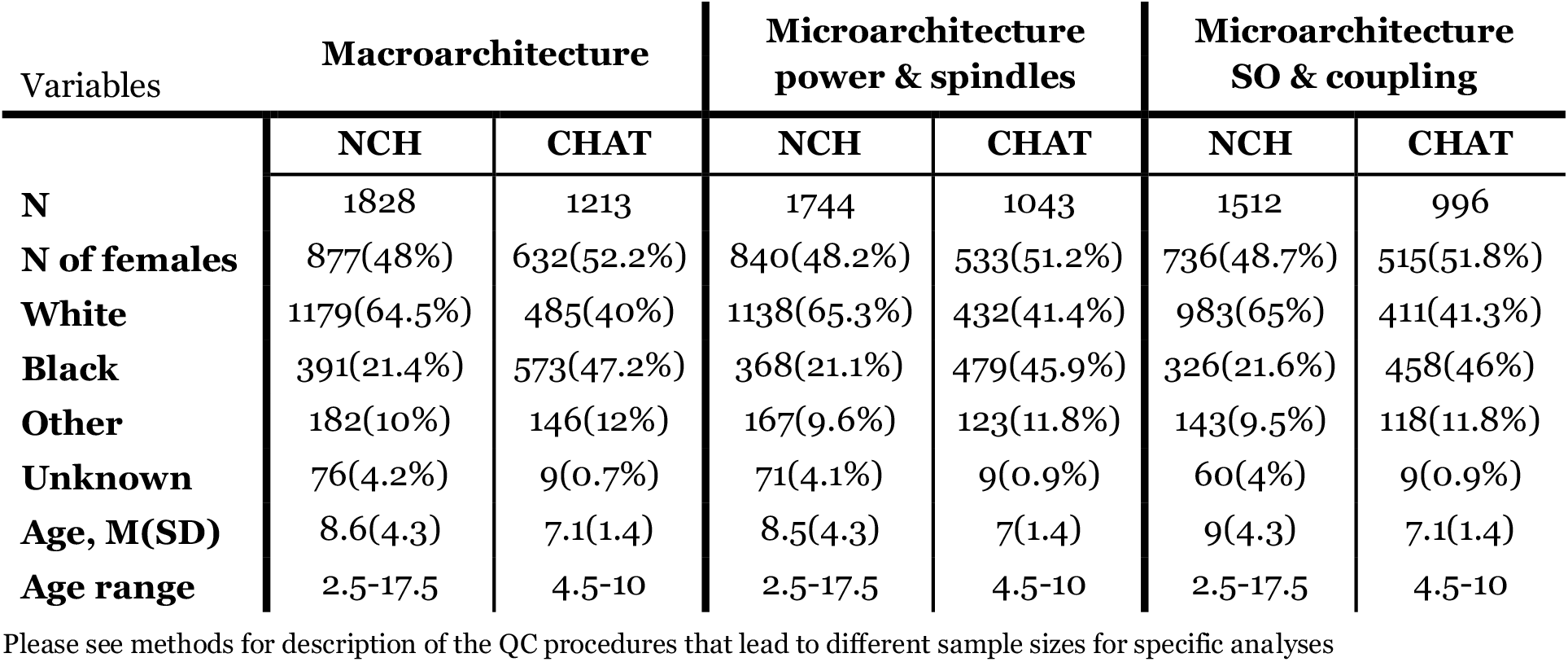
Demographics characteristics (excluding NDD groups)

**Table 2.** Distribution of Gender and Age by NDD group and Availability of Sleep Data in the NCH cohort.

| Disorders | Macroarchitecture |  | Microarchitecture power & spindles |  | Microarchitecture SO & coupling |  |
| --- | --- | --- | --- | --- | --- | --- |
|  | N (N of females) | Age, M(SD) range | N (N of females) | Age, M(SD) range | N (N of females) | Age, M(SD) range |
| **ASD** | 196 (45) | 9.5 (4.3)*<br>2.5-17.4 | 169 (39) | 9.3 (4.3)*<br>2.5-17.4 | 141 (30) | 9.6 (4.3)<br>2.7-17.1 |
| **ADHD** | 525 (149) | 10.9 (3.5)*<br>2.6-17.5 | 483 (136) | 10.9 (3.5)*<br>2.6-17.5 | 428 (119) | 11.2 (3.5)*<br>2.6-17.5 |
| **Intellectual disability** | 167 (60) | 10.9 (3.7)*<br>2.9-17.4 | 122 (45) | 11 (3.6)*<br>2.9-17.4 | 95 (33) | 11.3 (3.6)*<br>2.9-17.1 |
| **Down Syndrome** | 140 (56) | 8.1 (4.4)<br>2.5-17.4 | 122 (49) | 8.2 (4.5)<br>2.5-17.4 | 70 (23) | 8.7 (4.7)<br>2.5-17.3 |
| **Cerebral Palsy** | 138 (60) | 7.5 (3.8)*<br>2.6-17 | 93 (48) | 7.6 (3.8)*<br>2.6-17 | 79 (41) | 8 (3.8)*<br>2.6-17 |
| **Epilepsy** | 242 (104) | 8.7 (4)<br>2.5-17.4 | 179 (84) | 9 (3.8)<br>2.5-17.4 | 151 (72) | 9.2 (3.8)<br>2.6-17.4 |

\* - $p < 0.05$ : mean age is significantly different compared to the rest of the sample

### Sleep macro-architecture in children without NDD

Macro-architecture metrics (i.e. those derived from the hypnogram based on manual staging) showed marked age-related changes. Congruently in both samples, total sleep time (TST) exhibited a marked linear reduction with age (controlling for race and sex) in both NCH (*r* = -0.26, *p* < 10^−15^) and CHAT (*r* = -0.13, *p* = 9×10^−6^), as did sleep maintenance efficiency (SME) (*r* = -0.17, *p* = 2×10^−12^ in NCH, *r* = -0.07, *p* = 0.021 in CHAT) **(Figure 1A)**. Note that age-related effect sizes in CHAT are expected to be attenuated and/or more variable than in NCH, due to the narrower age range. Sleep also grew more fragmented with age in both samples, based on increases in the sleep fragmentation index (SFI) and duration of wake after sleep onset (WASO). The macro-architectural feature showing the strongest age-related change in both datasets was the number of NR sleep cycles (*r* = -0.38, *p* < 10^−15^ in NCH and *r* =-0.13, p *=* 4×10^−6^ in CHAT), which remained significant even after covarying for TST (p < 10^−15^ in NCH and p=0.002 in CHAT). At the same time, the average duration of sleep cycles increased with age in both samples, although the number of cycles was still the greatest determinant of TST (e.g. in NCH, *r* = 0.50 for cycle number compared to *r* = 0.12 for cycle duration).

**Figure 1.**
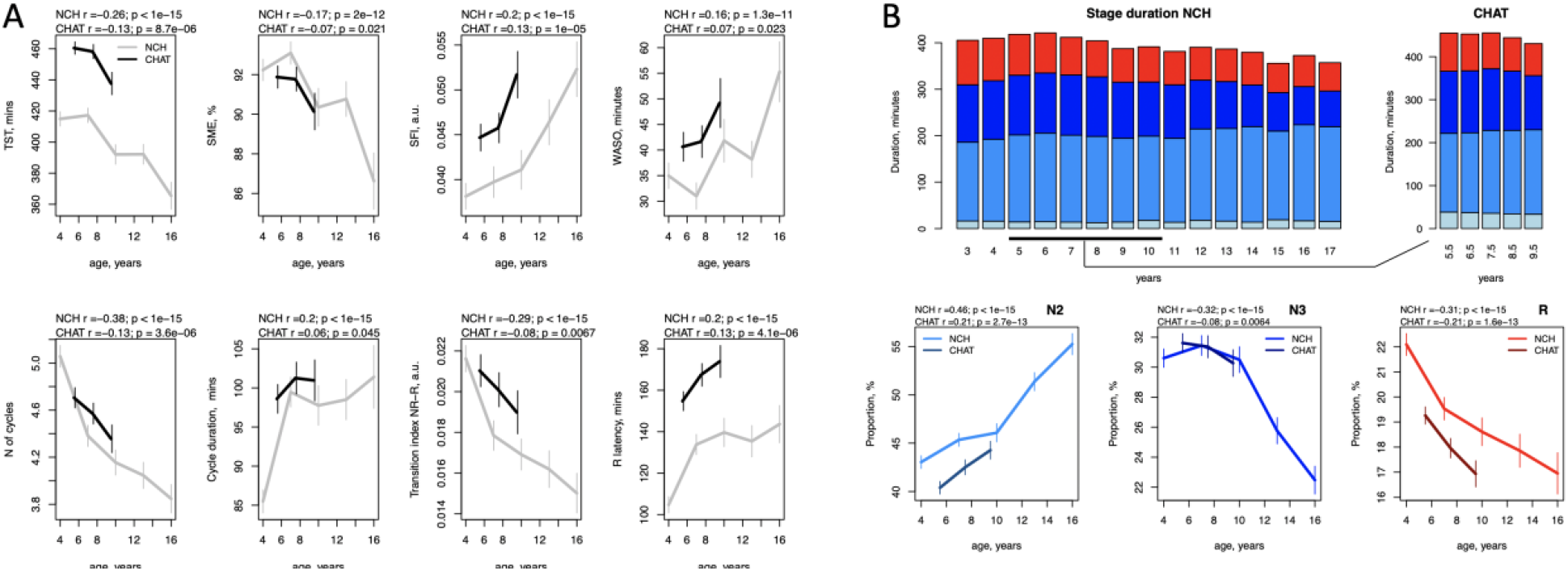
Development of sleep macro-architecture. A – Sleep macro-metrics stratified by age (3-year nonoverlapping windows from 2.5 to 17.5 years in NCH databank and 2-year nonoverlapping windows from 4.5 to 10 years in CHAT). The error bars represent 95% confidence intervals. B. Top row represents duration of sleep stages in minutes from 3 to 16 years (N1 – light blue, N2 – blue, N3 – dark blue, R – red); second row as in A, for N2, N3, R proportion with respect to total sleep time. Total sleep time (TST), sleep maintenance efficiency (SME), sleep fragmentation index (SFI), wake time duration after sleep onset (WASO), transition index between NR and R sleep (TI NR-R).

Despite statistically consistent associations with age, the age-adjusted absolute values of many macro-architectural metrics varied greatly between NCH and CHAT, a pattern observed in other contexts (e.g. comparing stage duration statistics between two elderly PSG cohorts, Djonlagic et al., 2021). Although such cohort effects may primarily reflect different recording contexts or other technical factors, rather than physiological differences between these populations, inasmuch as macro-architectural measures are susceptible to cohort-specific measurement biases, these issues may present challenges for the transferability of predictive models based on macro-architectural metrics.

As others have reported (Baker et al., 2016; Feinberg et al., 2012), sleep stage composition changed profoundly across this developmental period (**Figure 1B**). We observed similar effects in both cohorts, with one exception for N1 (**Figure 1B, top row**). N1 duration also showed the largest absolute difference between the datasets, possibly reflecting variations in manual staging protocols and the intrinsically low construct validity of N1 as a distinct and atomic physiological state, also echoing Djonlagic et al. (2021).

Stage N2 duration increased with age from 177 to 201 minutes between 4 and 16 years of age in NCH sample (*p* <10^−13^ in NCH, *p* = 0.003 in CHAT). In contrast, N3 sleep reduced with age from 126 minutes in 4-year-olds to 81 minutes in 16-year-olds (p < 10^−15^ in NCH and p = 10^−5^ in CHAT) suggesting an age-related reduction in mean NR depth. In general, the proportion of time spent in all NR (stages N1, N2 and N3) increased with age (*r* = 0.29, *p* < 10^−15^ in NCH and *r* = 0.19, *p* = 3×10^−10^ in CHAT) while stage R displayed age-associated reduction in both duration and proportion (*p*<10^−15^ in NCH and *p*<10^−13^ in CHAT) from 92 to 64 minutes between 4 and 16 years.

Further, stage R latency increased with age (*r* = 0.2, *p* < 10^−15^ in NCH, *r* = 0.13, *p*=7×10^−6^ in CHAT), although this effect was not significant after controlling for R duration and mean sleep cycle duration (data not shown). There were also fewer transitions between NR and R periods with age in both datasets.

### Sleep EEG spectral characteristics

Within sleep stage, spectral power across classical frequency bands displayed pronounced age-dependent changes (**Figure 2**), most notably a reduction in absolute spectral power in slower frequency bands (e.g. the largest effect size is illustrated **in Supp. Figure 2 A** – delta band absolute power during stage R, *r* = -0.82 in NCH and *r* = -0.3 in CHAT). However, these effects likely reflect gross changes in the amplitude of the sleep EEG (most evident for slower bands that have higher power due to the 1/*f* nature of the power spectrum); that they were observed uniformly across sleep stages and channels suggests that these effects may not be specific to sleep neurophysiology (versus gross anatomical changes, for example).

**Figure 2.**
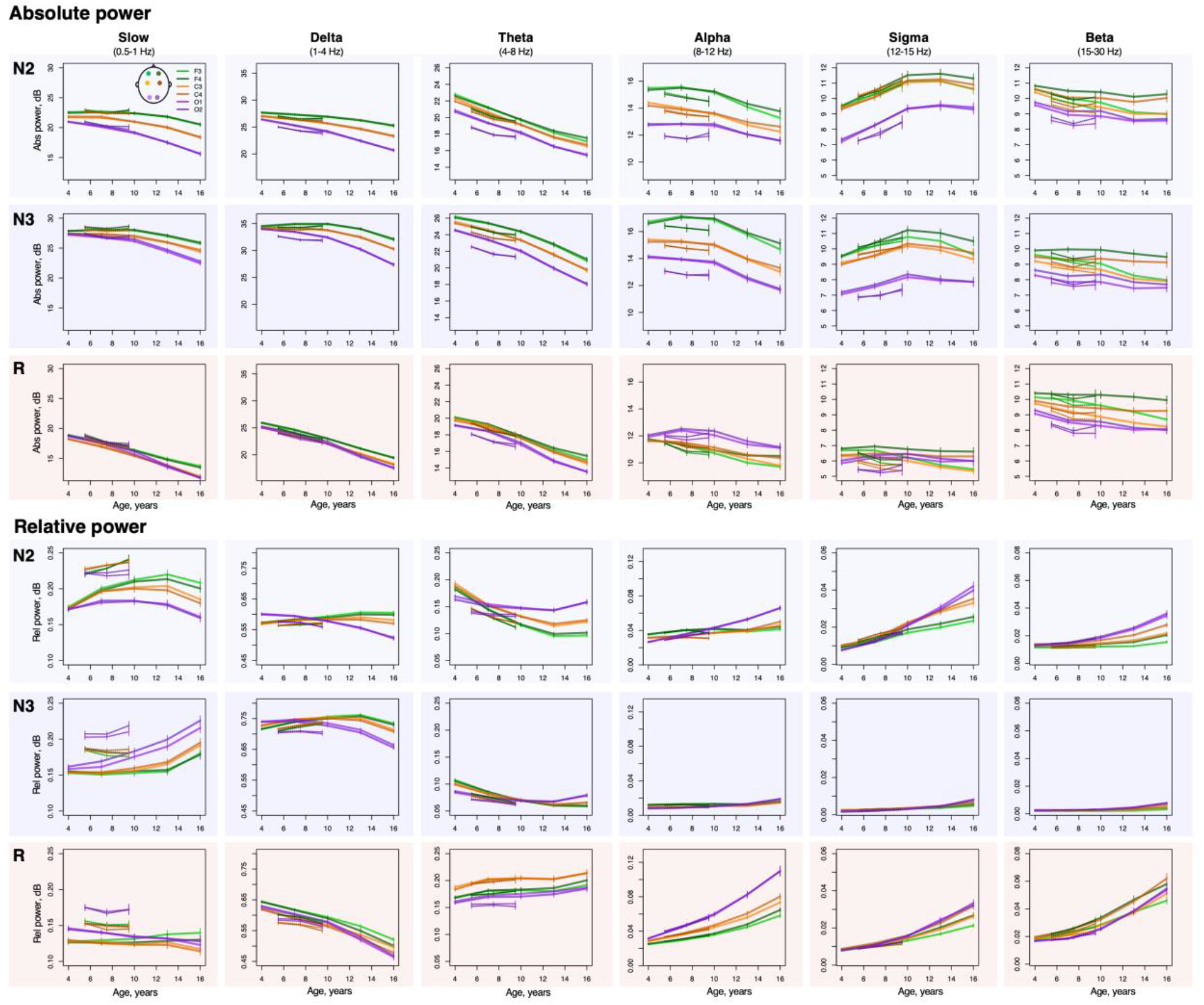
Developmental changes in spectral power. Developmental trajectories with 95% confidence intervals produced by averaging sleep estimates across individuals within 3-year nonoverlapping windows from 2.5 to 17.5 years in NCH databank and 2-year nonoverlapping windows from 4.5 to 10 years in CHAT) for absolute (top three rows) and relative (bottom three rows) across sleep stages (N2, N3, R, rows), bands (columns) and channels.

In contrast, absolute sigma power increased with age most strongly during N2 (*p* <10^−15^ in NCH and *p* < 0.001 in CHAT across all channels with strongest effects at O1, *r* = 0.39 and *r* = 0.16 for NCH and CHAT respectively). A smaller but still significant increase was observed during N3, but not REM. Modelling with higher-order age terms suggested a nonlinear trajectory (based on Akaike’s and Bayesian information criteria), reflecting a slight decline in power occurring in adolescence (**Figure 2**). Relative sigma power (normalized by total power 0.5 - 50 Hz to account for age-related changes in total power) similarly showed the strongest age-related changes (e.g. during N2 at O1, *r* = 0.76 in NCH and *r* = 0.4 in CHAT, **Supp. Figure 2 B**, with similar effects in N3, all *p* <10^−15^ in NCH and p < 10^−5^ in CHAT).

Other frequency bands and stages showed marked developmental changes in relative power with qualitatively distinct stage- and topographically specific developmental trajectories (**Figure 2**). Considering NCH only, delta power decreased with age during R across all channels (all *p* < 10^−15^), but increased with age in frontal channels, during N2/N3, with a peak around the age of puberty. In occipital channels, delta power decreased with age across all stages (all *p* < 10^−15^). In contrast, theta power increased with age during R (all *p* <10^−10^) but decreased with age during N2/N3 (all *p* <10^−15^) in frontal and central channels. For the comparable age range, we observed broadly consistent patterns of NR age-related change in CHAT. Finally, during REM, relative alpha, sigma and beta power increased with age (all *p* < 10^−15^ in NCH and all *p* < 10^−10^ in CHAT).

### Spindles, slow oscillations and their coupling

Across childhood and adolescence, we observed multiple changes in NR sleep spindles (**Figure 3**). Although the density (count per minute) of both slow and fast spindles (SS and FS respectively, targeting 11 Hz and 15 Hz activity, with approximately +/-2 Hz around each central frequency) increased with age in frontal channels (all *p* < 10^−15^ in NCH and *p* < 10^−14^ in CHAT), their precise developmental trajectories were distinct (**Figure 3**). Whereas FS density linearly increased across all channels (all *p* < 10^−15^ in NCH and *p* < 10^−10^ in CHAT) from 0.9 spindles per minute at age of 4 to 1.9 at age of 16 at C3, SS density displayed an inverted-U profile, most pronounced in frontal channels and peaking around 10 years with 2.4 spindles per minute at F3 (compared to 1.5 and 1.9 spindles at 4 and 16 years of age, respectively).

**Figure 3.**
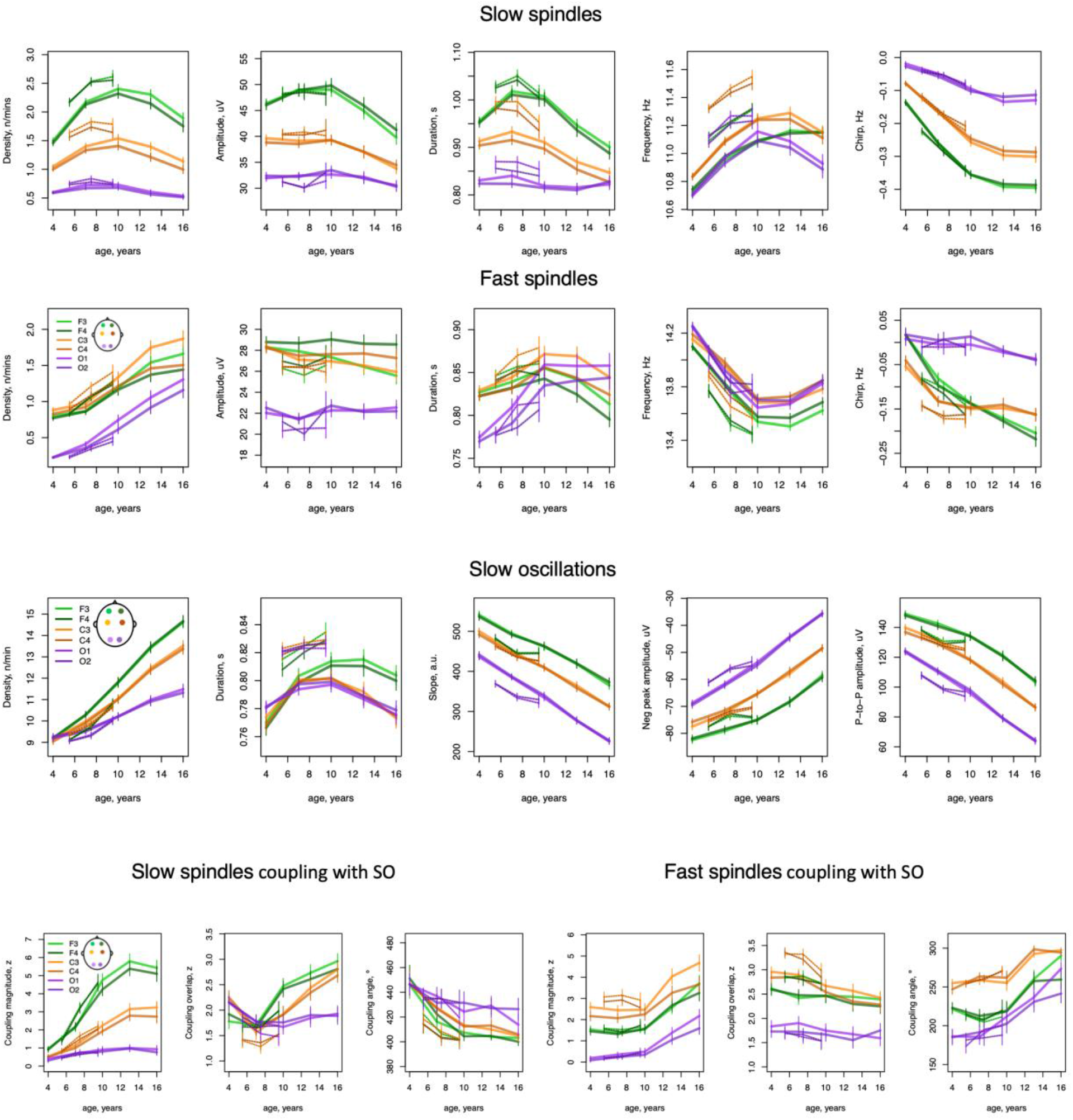
Developmental trajectories of spindles, slow oscillation, and their coupling. Developmental curves with 95% confidence intervals generated by averaging sleep estimates within 3-year nonoverlapping windows from 2.5 to 17.5 years in NCH databank and 2-year nonoverlapping windows from 4.5 to 10 years in CHAT) for slow (top row) and fast (second row) spindle characteristics, SO parameters (third row) and coupling between SS/FS and SO (bottom row) during N2 across all channels.

Spindle morphology, not rate of occurrence, showed the most marked age-related changes, however. In particular, intra-spindle deceleration (sometimes called “chirp”, a typical characteristic of both fast and slow spindles) grew more pronounced with age, especially for SS (**Figure 3**, *p* across all channels < 10^−15^ in NCH and *p* < 10^−7^ in CHAT, with strongest effects at F3: *r* = -0.66 and *r* = -0.38 in NCH and CHAT respectively (**Sup. Figure 2 C**) as opposed to largest effect in FS density *r* = 0.53 and *r* = 0.29 in NCH and CHAT).

Average spindle frequency varied markedly with increasing age, but differently for slow and fast spindles **(Figure 3)**: whereas SS frequency became faster with age (across all channels *p* < 10^−15^ in NCH and *p* < 10^−10^ in CHAT), FS frequency slowed down (all *p*<10^−15^ in NCH and *p*<10^−8^ in CHAT). Opposing directions of change in frequency of SS and FS was also observed in older adults (Djonlagic et al., 2020). Notably, both SS and FS developmental trajectories were highly non-linear with SS maximum and FS minimum frequency observed around 13 years of age.

We detected slow oscillations (SOs) by identifying zero-crossings in 0.5-4 Hz bandpass-filtered signals and applying fixed duration but relative amplitude thresholds (see Methods for details). As for spindles, SO density increased dramatically with age (**Figure 3**) across all channels in both cohorts (all *p* < 10^−15^ in NCH and *p* < 10^−15^ in CHAT). The steepest increase observed was at F4 with 10 SOs per minute at age of 4 and 15 at 16 years of age (*r* = 0.7 in NCH and *r* = 0.38 in CHAT, **Sup. Figure 2D**).

With respect to SO morphology, average negative peak amplitude, peak-to-peak amplitude, and the slope between negative and positive peaks decreased linearly across all channels (p < 10^−15^ in NCH and *p* < 10^−5^ in CHAT, with the largest effect for peak-to-peak amplitude at O1, *r* = -0.74 in NCH and *r* = -0.23 in CHAT, **Sup. Figure 2F**). SO duration was the only tested parameter to express a marked non-linear trajectory, with a steep increase from 4 to 10 years and a steady decline after age 10-12 years (**Figure 3**).

SO properties are necessarily dependent upon the criteria used to detect them. In our primary analyses, based on optimizing the strength of observed spindle/SO coupling, we elected to use a relative amplitude threshold (see Methods for details). For example, 63% of individuals had significant FS/SO overlap at C3 in NCH (66% in CHAT) versus 58% and 56% when an absolute SO amplitude threshold was used. The choice of SO detection criteria impacted patterns of age-related change, however. Indeed, using an absolute threshold, age-related trends for SO rate, slope and amplitude were reversed – the former decreased with age while the latter two increased (**Sup. Figure 3**). This apparent contradiction reflects a general decrease in SO activity with age, which is congruent with the observed age-related decrease in slow and delta band power. Reduced SO activity consequently lowers any relative threshold, which can in turn lead to relatively more events being detected. The optimal choice of threshold is an empirical question that will depend on the subsequent analyses, which underscores the importance of always explicitly reporting the type of detection thresholds used.

Returning to the original relative-threshold set of SO, we assessed spindle/SO coupling in three ways: 1) the tendency for spindles to preferentially occur non-uniformly with respect to SO phase (coupling magnitude), 2) the preferred SO phase at spindle peaks (coupling angle), and 3) the extent of any above-chance overlap between spindle and SO events, ignoring SO phase (coupling overlap). In general, SS showed more marked age-related changes, compared to FS (**Figure 3**). SS coupling increased in frontal and central channels in both cohorts (all *p* < 10^−15^ in NCH and *p* < 10^−1^ in CHAT: e.g. at F3 *r* = 0.56 in NCH and *r* = 0.44 in CHAT, **Supp. Figure 2E**).

With respect to FS coupling, an age-related increase was observed only in the NCH sample (across all channels *p* < 10^−15^), primarily driven by a steep increase in adolescence (**Figure 3**). The lack of CHAT replication here likely reflects the restricted age range: indeed, among NCH individuals 10 or under associations were greatly attenuated, compared to the same tests in the older NCH subsample (data not shown).

Next, we found that individual’s preferential SO phase angle at spindle peak (circular mean) shifted across development. In both cohorts, FS tended to occur before the SO positive peak, whereas SS tended to occur after it. However, with increasing age, both SS and FS shifted closer to the SO positive peak. Again, the most rapid change in preferred FS SO phase was during adolescence.

Finally, rates of above-chance gross overlap between SS and SOs increased with age at frontal channels (F3/F4 *p* < 10^−15^ in NCH and *p* < 0.05 in CHAT) whereas FS overlap showed a modest (albeit significant) decrease at central channels (C3/C4 *p* < 0.01 in NCH and *p* < 0.05 in CHAT). Unlike other SO metrics, age-related changes in coupling were generally similar despite different approaches for SO detection (**Sup. Figure 3**).

Given that the majority of individuals in both cohorts had sleep apnea and/or snoring, we additionally tested all sleep variables association with age including apnea-hypopnea index (AHI) as a covariate. The results were effectively identical after controlling for AHI with Pearson’s correlation > 0.99 between signed log-transformed original and AHI-controlled *p*-values, and comparable levels of significance.

### Brain age prediction using sleep macro- and micro-architecture measures

Above, we demonstrated 1) that age was very strongly associated with multiple sleep macro- and micro-architecture metrics, and 2) that findings were congruent for two samples from distinct (clinical versus research) sources. We next aimed to condense these multivariate developmental patterns into a single model to estimate chronological age from the sleep EEG, and then to test whether its deviation from observed age - i.e. *brain age gap* - could identify pathological neurodevelopment. We fit a simple linear regression of age on sleep macro and microarchitecture metrics adjusting for sex and race, using 70% of the NCH sample (after first excluding clinical subgroups). The remaining 30% was used as an initial testing set. As well as using CHAT as a second independent test dataset, we additionally utilized a new pediatric dataset with a wider age range (Pediatric Adenotonsillectomy Trial for Snoring, PATS: *N* = 627 [307 females], mean age 6.3 years, range 3 to 13 years). PATS was comprised of children with snoring from diverse ethnic backgrounds participating in a clinical trial; we used PATS as a third independent test sample in order to further assess the transferability of the NCH-trained model.

In all three test samples, the model predicted age with relatively high accuracy, indicating a high degree of transferability (**Figure 4A**). The highest performance was observed in the NCH testing set, where predicted and observed age correlated *r* = 0.95. Although the correlation was lower in CHAT and PATS datasets (*r* = 0.65 in CHAT, *r* = 0.88 in PATS), likely in part due to the narrower age ranges, the mean absolute error (MAE) in those samples was comparable with the NCH (MAE = 1.05 - 1.14 years). If taken separately, models based on only micro-architecture or spectral power metrics performed almost as well as the model using all metrics combined, whereas models based on only macro-architecture metrics led to the lowest accuracy of prediction (MAE > 1.5 years in all datasets) and much lower correlations (*r* = 0.56 in NCH, *r* = 0.22 in CHAT, *r* = 0.35 in PATS). Prediction accuracy was similar for boys and girls in all three testing samples (*p* > 0.05) even when participant sex was not included in the model. Nonetheless there were still significant (albeit relatively subtle) sex differences and sex-by-age interactions in multiple measures of sleep macro and microarchitecture consistent with the previous literature (Baker et al., 2020; Campbell et al., 2012) (**Sup. Figure 4**). The brain age gap (MAE and ME) was also not significantly associated (*p* > 0.05) with AHI in either NCH or CHAT.

**Figure 4.**
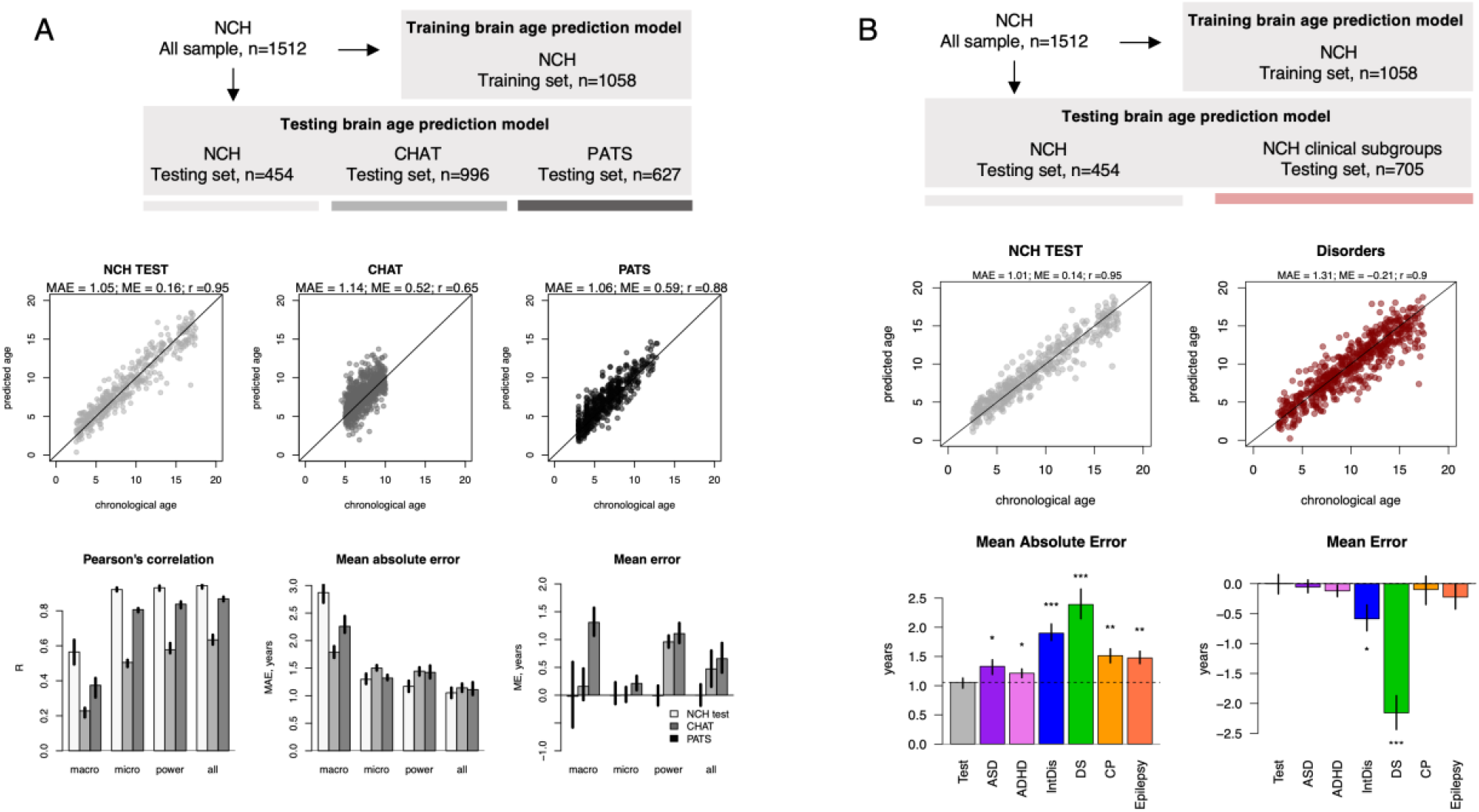
Brain age prediction based on sleep micro and macro-architecture in typical and clinical samples. First row describes training and testing sets and their sample sizes for age prediction in typical and clinical samples. Second row scatterplots show predicted vs true age for all testing sets. Bar plots in the third row (A) illustrate Pearson’s correlation (r), mean absolute error (MAE) and mean error (ME) between true and predicted age when prediction model used features taken from different domains of sleep metrics (macro, micro, spectral power and all combined). To assess how variable these estimates were, we repeated the analysis 50 times with different non-overlapping individuals from NCH sample randomly assigned to testing and training set each round (CHAT and PATS sample always stayed the same). Resulted 50 estimates of r, MAE, ME for each testing set were averaged and plotted in bar plots with error bars illustrating min and max estimates of those metrics across 50 rounds. Bar plots in the third row (B) illustrate MAE and ME for each clinical subgroup in comparison to the testing set. Similar to (A) we estimated variability of these estimates by repeating the analysis 50 times where training and testing set was changing each round, but the disorder subgroups remained unchanged. The resulting 50 estimates of MAE and ME were averaged and illustrated in bar plots (error bars are min and max values across 50 rounds). Stars indicate statistical difference between estimates in each clinical subgroup vs testing set (* - p <0.05, ** - p <0.01, *** - p <0.0 01).

### Brain age prediction and NDD

Brain age gap was significantly negative (i.e. younger than expected) in individuals with DS (ME = -2.2 years, p = 4×10^−11^) or intellectual disability (ME = -0.59 years, p = 0.043) subgroups (**Figure 6 B**). Interestingly, for DS the brain age gap effect was exacerbated with increasing age, suggesting greater developmental delays in older patients (**Figure 5 E**). Given prior reports of accelerated ageing (i.e. positive age gap) in DS adults based on structural MRI and epigenetic markers (Cole et al., 2017; Horvath et al., 2015), we sought to contextualize our finding of delayed development in DS, especially given that the NCH training sample was obligatorily limited to children and adolescents, combined with the fact that some sleep EEG metrics follow nonlinear, inverted-U trajectories across the lifespan (e.g. fast spindle density increases across childhood, peaks around 20 years of age and slowly declines thereafter, **Figure 5B**).

**Figure 5.**
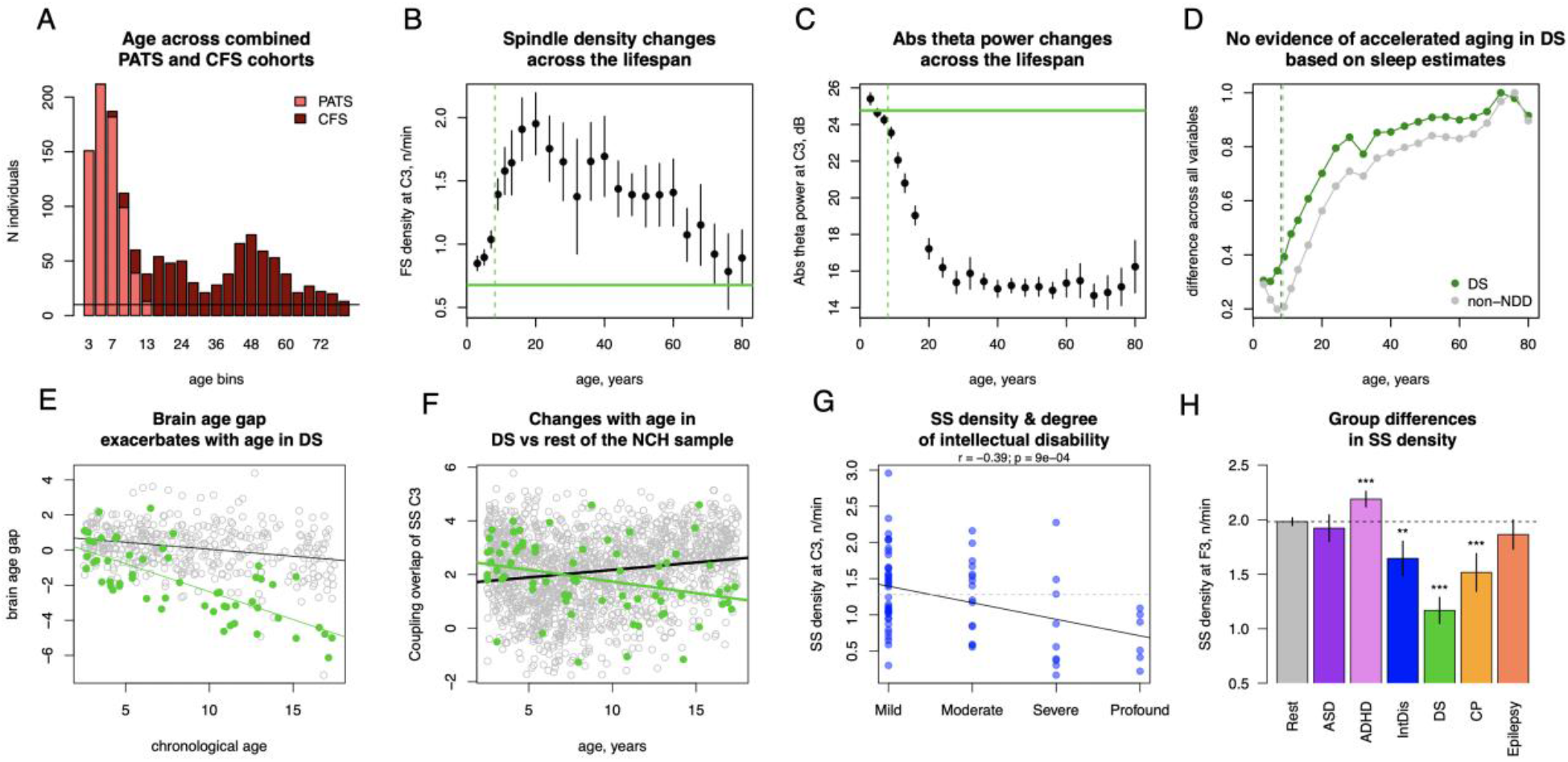
Examples of NDD effects on sleep micro-architecture. A – age distribution in the combined sample of PATS and CFS cohorts; B, C – fast spindle density and absolute theta power averaged across age bins (lines indicate 95% CI) in the combined sample in reference to the mean fast spindle density and absolute theta power across individuals with DS (green horizontal line, dashed vertical line illustrates mean age in DS subgroup; D – normalized mean absolute difference between DS group and each age bin in the combined sample across sleep macro- and microarchitecture estimates (in green, grey point indicate the same but for non-NDD sample). Vertical lines illustrate average age of DS and non-NDD groups. E – scatterplot illustrates changes in brain age gap with age in DS subgroup (in green) and the test sample. F – distinct developmental trajectory of SS coupling overlap at C3 in DS subgroup (in green) compared to the rest of non-NDD sample; G – correlation between SS density at C3 and degree of intellectual disability (grey dashed line illustrates the mean SS density at C3 for the non-NDD sample); H – SS density across subgroups and the rest of the sample at F3 (** - p < 0.01, *** - p < 0.001).

We therefore estimated the developmental trajectories of sleep EEG metrics in new sample spanning both childhood and adulthood: PATS combined with the Cleveland Family Study (*N*=730, 401 females, mean age of 41 spanning from 6.8 to 89 years) cohorts (**Figure 5A**). As expected, one can identify a group of older adults with numerically equivalent mean spindle densities compared to young children, including the DS group (**Figure 5B**). However, when looking across all other sleep metrics, the DS children group in no way resemble the older adult group, i.e. as one might expect under the hypothesis of extreme accelerated aging, either for individuals metrics such as absolute theta power (**Figure 5C**) or when considering the similarity across a composite of sleep metrics (**Figure 5D**). That is, when considering the broad profile of all age-dependent sleep metrics, the DS group (mean age of 8 years) was the most similar to the average for children aged 6 years (i.e. consistent with a delay) rather than any older adolescent or adult group.

In absolute terms, the brain age gap was significantly larger across all NDD subgroups: DS (MAE = 2.4 years, p = 2×10^−7^), intellectual disability (MAE = 1.9 years, p = 9×10^−6^), CP (MAE = 1.5 years, p = 0.008), epilepsy (MAE = 1.5 years, p = 0.003), ASD (MAE = 1.33 years, p = 0.04) and ADHD (MAE = 1.21 years, p = 0.03) groups, consistent with greater levels of heterogeneity among NDD groups. (The mean MAE across typical CHAT and PATS samples was consistently below 1.18 years.)

To gain some insight into which sleep alterations were driving these brain age discrepancies in NDD, we performed a series of exploratory analyses comparing each NDD subgroup with the rest of the non-NDD NCH sample, statistically controlling for i) the other, potentially overlapping NDD subgroups, ii) diagnoses of major sleep disorders, iii) contemporaneous use of medications likely to impact sleep, iv) age, v) sex, vi) race and vii) AHI (see Methods for details).

Individuals with DS – the subgroup with the largest mean discrepancy between predicted and chronological age – showed the largest number of significant group differences and age-by-group interactions (**Sup. Figure 5**). The largest effect sizes were for spindle characteristics – lower SS spindle density (1.1 n/min in DS vs 1.9 in the non-NDD sample at F4, *p* < 10^−15^), duration (at F3 *b*_*st*_ = -1.1, *p* < 10^−15^), less pronounced chirp (at C3 *b*_*st*_ = 0.86, *p* < 10^−15^) and absolute beta power (at F3 during N2 *b*_*st*_ = 0.94, *p* < 10^−15^, **Supp. Tables 3-5**). In terms of age-related trajectories, coupling between SO and spindle metrics were among those showing stronger age-by-group interactions. For example, coupling overlap between SO and SS at F3 significantly decreased with age in DS (**Figure 5F**, *r* = -0.28, *p* = 0.02) but increased in the non-NDD sample (*r* = 0.17, *p* < 10^−10^).

Despite significant deviations in both MAE and ME, the number of altered sleep estimates were much lower in the intellectual disability group compared to the DS group; these were largely related to spindle frequency characteristics (**Supp Figure 5**). SS density was associated with the degree of intellectual disability (the largest effect at C3 *r* = -0.39, *p* = 0.0009, **Figure 5G**) and was in general decreased compared to individuals without NDD in frontal channels (the largest effect at F3 *b*_*st*_ = -0.29, *p* = 0.003, **Supp. Tables 5, Fig. 5H**).

Finally, multiple nominally significant group and age-by-group alterations were detected for ADHD, CP and Epilepsy (Supp. **Figure 5, Sup. Table 3-6** contain full reports of uncorrected results). Except for ASD, all NDD-subgroups had common significant (*p* < 0.05) deficit of stage R sleep expressed as reduced duration (in DS), proportion (in intellectual disabilities) or both (in CP and epilepsy) and stage R latency (in ADHD) even after adjusting for sleep disorders and relevant medication.

## Discussion

Using both clinical and research datasets, we identified consistent patterns of age-related change during childhood and adolescence, for multiple facets of sleep macro- and micro-architecture. We further showed that, when combined, the different sleep EEG metrics we examined could reliably predict an individual’s age in pediatric populations, and that the resulting models were broadly transferable across different cohorts. Finally, we showed that some NDD subgroups (primarily DS) exhibited systematic differences in their predictions of age, reflecting multiple disruptions of sleep architecture.

### Macro-architecture changes with age

Confirming numerous previous reports of decreasing total sleep time during the first two decades of life (Galland et al., 2012; Iglowstein et al., 2003; Ohayon et al., 2004), we additionally showed significant changes in other sleep macro-architecture metrics. Congruently with a large cross-sectional study of children without sleep complaints (Scholle et al., 2011), we found an increase in sleep cycle duration, stage R latency and a decrease in a number of sleep cycles with age. With respect to sleep efficiency, for which both age-related increases (Baker et al., 2012; Scholle et al., 2011) as well as decreases (Baker et al., 2016) have been reported, our findings pointed to a significant decline in both cohorts.

Major developmental changes were also observed with respect to sleep stage composition. Prior reports of developmental trajectories of NR sleep showed consistent findings of an increase in N2 and a reduction in N3 with age (Baker et al., 2016; Jenni & Carskadon, 2004; Ohayon et al., 2004; Scholle et al., 2011; Tarokh et al., 2011; Tarokh & Carskadon, 2010), as found in our analyses. In contrast, the reports diverge for stage R sleep showing either an increase (Baker et al., 2016; Feinberg et al., 2012; Ohayon et al., 2004), no change (Tarokh et al., 2011; Tarokh & Carskadon, 2010), or a decrease (Scholle et al., 2011) in stage R percentage and/or duration. Our analysis -- which represents the largest study to date -- supports a decrease in R sleep across ages 2.5 to 17.5 years.

### EEG spectral composition changes with age

Multiple studies have reported age-related changes in absolute delta power during NR sleep, suggesting an inverted-U trajectory: an increase from childhood to adolescence (Feinberg et al., 2012) and reduction from adolescence to adulthood (Baker et al., 2012, 2016; Feinberg & Campbell, 2013; Jenni & Carskadon, 2004; Tarokh & Carskadon, 2010). During R sleep, a linear decrease in delta power has also been documented (Feinberg & Campbell, 2013). Theta power has also been reported to decline with age across all sleep stages (Feinberg & Campbell, 2013; Gaudreau et al., 2001).

We confirmed topographical patterns of relative delta power change across a large and diverse sample. In a sample of 55 individuals (2.4-19.4 years), the scalp location with maximal SWA (1-4.5 Hz) power shifted with age in a posterior-to-anterior direction (Kurth, Ringli, et al., 2010), mirroring the pattern of cortical thinning during childhood and adolescence. Several studies utilizing both MRI and sleep EEG reported a link between slow-wave activity during NR sleep and cortical thickness and/or gray matter volume (Buchmann et al., 2011; Goldstone et al., 2018). In general, the developmental decline in total power, as well as in delta and theta frequency bands, is not specific to sleep but also widely reported in wake (Barriga-Paulino et al., 2011; Boord et al., 2007; Gasser et al., 1988; Whitford et al., 2006), and has been similarly linked to reduction in cortical gray matter volume, and cerebral metabolic rate (Boord et al., 2007; Whitford et al., 2006). Thus, similar developmental changes in EEG spectral composition during both wake and sleep might reflect global structural changes in the brain, such as synaptic pruning that underlies normal cortical maturation and are not state-specific (Segalowitz et al., 2010).

In contrast to the general decline in total power, absolute sigma power increased with age. A similar increase was previously reported from 10 to 12 years in a longitudinal sample (Tarokh & Carskadon, 2010) but there are also reports of higher absolute sigma in children compared to adults (Gaudreau et al., 2001). Another longitudinal sample reported a complex trajectory of sigma power where it increased linearly from 6 to 12 years of age and then decreased from 12 to 16 (Feinberg & Campbell, 2013; Kurth, Ringli, et al., 2010), matching our observations of inverted-U trajectories in spindle activity.

When normalized by total power, sigma power displayed one of the strongest age-related effects. During NR sleep, this increase can be attributed to spindle maturation (discussed below). During R sleep, relative sigma power age-related increases were also accompanied by increases in relative alpha and beta power. While sleep studies reporting age-related changes in relative power are scarce, similar findings of an overall increase in higher frequencies are well reported during wake (Dustman et al., 1999; Gasser et al., 1988; Soroko et al., 2014), once again pointing to brain maturation processes evident across both sleep and wake.

Although, to our knowledge, there are no prior studies explicitly reporting age-related changes in relative power across classical frequency bands in sleep, our results indicate that relative power is a good metric to highlight topographical-, stage- and frequency-dependent aspects of developmental changes in the sleep EEG. For example, we observed distinct and sometimes contrasting trajectories between occipital and anterior channels as well as between NR and R sleep stages (e.g. in delta and theta frequency bands), that can be practically leveraged when developing multi-channel tools for automatic stage classification in children and adolescents.

### NREM microarchitecture

Rapid increases in spindle density during childhood have been linked to maturation of thalamocortical circuits (Fernandez & Lüthi, 2020). Our results tend to confirm previous reports of slow spindle density increasing during childhood and reaching its peak around puberty (Purcell et al., 2017; Scholle et al., 2007). SS duration also was shown to exhibit a similar developmental pattern, peaking slightly earlier than density, confirming previous reports (Scholle et al., 2007). In contrast, FS density increased through adolescence, congruent with recent studies (Goldstone et al., 2019; Purcell et al., 2017).

One of the most frequently reported findings is that the spindle frequency peak estimated from examination of the power spectrum increases with age (Campbell & Feinberg, 2016; Hahn et al., 2020; Tarokh et al., 2011). Although one early study showed that this is true for both frontal and centro-parietal spindles (Shinomiya et al., 1999), tracking the spindle frequency based on the sigma peak for both slow and fast spindle can be problematic given that the sigma peak in children falls within a range of slow spindles (Hoedlmoser et al., 2014). Alternatively, sigma peak increase could potentially reflect a shift in the ratio of slow versus fast spindle density with fast spindles becoming more prevalent after puberty. Our findings were based on spindle detection as discrete events, an approach that allowed us to estimate change in frequency for both slow and fast spindles in a more direct way. We showed that while SS frequency increases with age congruently with previous reports based on the sigma power peak, the FS frequency decreases. Interestingly, such a pattern appears to be opposite of the aging effect in aging adults (50s to 80s) where the SS and FS frequencies tend to diverge with SS becoming slower and FS faster (Djonlagic et al., 2020). A recent study that studied spindles in three groups of adolescents found that both slow and fast spindles frequency was increased with age (Bocskai et al., 2022). However, in that study, slow and fast spindles were distinguished only using topographical differences (all spindles detected in frontal channels were declared slow and all spindles in centro-parietal channels fast).

We also report a previously undocumented finding for intra-spindle frequency change (or spindle “chirp”). This metric is usually negative, reflecting the characteristic deceleration of both SS and FS (Andrillon et al., 2011). Previous literature has suggested that chirp might be linked to spindle termination mechanisms and cortical modulation (Carvalho et al., 2014). We found that intra-spindle deceleration intensifies over childhood and adolescence, showing some of the most pronounced age-related changes in our study.

SOs also displayed some of the strongest age-related changes in our cohort. SO rate per minute increased dramatically from childhood to adolescence while SO amplitude and slope decreased. Age-related decreases in SO amplitude and slope were reported previously in a small (*N*=14) sample (Kurth, Jenni, et al., 2010). We also reported, however, that the interpretation of age-related trends is obligatorily highly dependent on the choice of the SO amplitude detection threshold. Prior work showed that coupling between SOs and spindles also increased from childhood to adolescence in longitudinal sample of 33 individuals (Hahn et al., 2020); this result was confirmed in our sample and extended to show that the strongest changes are for SS coupling magnitude.

### Estimating brain age from the sleep EEG

We showed that individual differences in sleep macro- and micro-architecture can be summarized using simple methods to generate a highly accurate predictor of chronological age that is transferable across multiple independent samples. As has been widely employed by many groups using different brain imaging modalities (as well as epigenetics and other biomarkers), the difference between predicted “biological” age and observed chronological age can be interpreted as a measure of development and health (Franke & Gaser, 2019). In pediatric populations, studies using structural MRI (Franke et al., 2012; Hong et al., 2020) have been able to predict age with high accuracy (*r* > 0.9). Age has also been predicted using functional MRI (Li et al., 2018; Lund et al., 2022; Qin et al., 2015), albeit with lower accuracy (*r* = 0.54 - 0.73). While a few studies using sleep EEG in adult cohorts achieve good results – *r* = 0.82-0.93 (Nygate et al., 2021; Sun et al., 2019), to our knowledge this is the first study to demonstrate this in a pediatric cohort.

One recent report used resting state EEG spectral power to estimate brain age in a cohort of 5-18 year-olds, with an average MAE of 1.2 years (Vandenbosch et al., 2019). This MAE is broadly similar to our results (MAE < 1.17 years), although we note that are our results are based on performance in two independent samples, not only by means of cross-validation within the same sample, as in (Vandenbosch et al., 2019). Given that many spectral age-related changes were evident in wake as well as sleep, we might expect similar performance for age prediction using either wake or sleep EEG. However, it remains an open and empirical question as to how highly correlated brain age estimates are when based on different modalities (MRI versus EEG) or different physiological states (wake versus sleep). Note that better prediction of chronological age is not, in itself, necessarily the most relevant factor: as a trivial conceptual example, a model that achieved perfect prediction (r=1, MAE=0) would be useless. Furthermore, even if two approaches have identical performance with respect to prediction of chronological age *per se*, they may still yield very different results with respect to how the model residuals (i.e. the so-called brain age gap) relate to brain development and health.

To address the question of the biologically or clinically relevant properties of the brain age gap, we estimated it in NDD subgroups as well as typically developing children. The largest deviations in both absolute and signed values (MAE and ME) were seen in individuals with DS and intellectual disability, with both groups showing negative gaps, consistent with delayed brain development. While we are not aware of other studies reporting on brain age in children with DS, a similar analysis was conducted for 46 adults (age range 28 - 65 years) with DS using structural MRI. In contrast to our results, they reported a positive brain age gap interpreted as accelerated aging (Cole et al., 2017), finding that it was related to increased beta amyloid deposition and cognitive decline. With the use of DNA methylation levels to calculate an ‘epigenetic clock,’ Horvath and colleagues also pointed to accelerating biological aging in brain and blood tissue (Horvath et al., 2015), with evidence that such advanced ageing of blood samples begins prenatally (Xu et al., 2022). Likewise, as well as shorter life expectancies generally, adults with DS display older biological age based on multiple physiological measures (e.g. BMI, blood pressure, etc) (Nakamura & Tanaka, 1998). When cognitive and behavioral levels were assessed, however, individuals with DS tend to have lower developmental age (Gameren-Oosterom et al., 2011), similar to our findings of children with DS being the most similar to younger age children with respect to their sleep macro and microarchitecture. Such results support the notion that accelerated/decelerated aging patterns are not universal and can be tissue and system-specific (Horvath et al., 2015), as well as that brain age based on the sleep EEG may be reflective of cognitive and behavioral development.

Additional analyses controlling for chronological age revealed alterations in multiple sleep macro and micro-architecture metrics in the DS subgroup, many of which were the opposite of typical age-related changes, suggesting altered developmental patterns in DS. For example, we saw a global increase in absolute spectral power in DS versus an age-related decrease in the control groups. Likewise, individuals with DS had reduced SS, FS and SO density across ages 4 to 16, counter to the marked age-related increases in these metrics in this age range. With the exception of one report of increased higher total spectral power (Sibarani et al., 2022), the results of which we confirm here, a fuller assessment of sleep microarchitecture in DS has not been conducted and so our findings provide an important developmental perspective on abnormalities associated with DS.

We also observed a consistently younger functional pattern in the sleeping brain in individuals with intellectual disabilities. In terms of sleep microarchitecture, spindle frequency metrics expressed the most marked alterations. While it is hard to compare our findings to the existing literature due to scarcity of reports available, two reports concluded that children with intellectual disabilities – especially those with more severe impairments – had decreased spindle density based on visual detection (Shibagaki et al., 1980; Shibagaki & Kiyono, 1983). Our findings also pointed to reduction in SS density in ID, that was more profound with the higher degree of intellectual disability.

While other NDD subgroups – ASD, ADHD, CP and epilepsy – did not express consistent shifts towards either younger or older brain (ME), they all expressed larger brain age gaps in absolute terms (MAE). This may indicate considerable heterogeneity within these disorders: indeed, this has been previously reported in other contexts for ASD and ADHD (Dajani et al., 2016; Jeste et al., 2015) as well as CP (Rosello et al., 2021) and epilepsy (Pack, 2019). Alternatively, these results could reflect group-level differences in the sleep EEG leading to increased noise in the age prediction model, as NDD groups were excluded from the primary NCH model fitting. In general, future studies will be needed to fully evaluate the relative merits of different brain age metrics, and to determine whether ones based on the sleep EEG offer additional, unique information or not, as well as how brain age alterations may vary over the course of a disease.

Except ASD, all subgroups displayed alterations in stage R sleep, either in terms of absolute duration, relative duration or stage R latency. This is consistent with previous reports of stage R deficits in DS (Spanò et al., 2018), CP (Hayashi et al., 1990), epilepsy (Sadak et al., 2022), and intellectual disability (Esposito & Carotenuto, 2014), supporting the notion that R deficits are common characteristics across NDDs. Previous studies reported that lower R duration was associated with worse cognitive performance and mortality in older individuals (Djonlagic et al., 2020; Leary et al., 2020). As such, R sleep metrics may not be good candidates for condition-specific biomarkers, but rather reflect pathophysiological alterations shared between distinct disorders.

### Caveats & conclusions

In summary, the present study provides a comprehensive assessment of age-related changes in sleep macro and microarchitecture, based on a large sample from multiple cohorts spanning the first two decades of life. Nonetheless, certain constraints should be mentioned. One obvious limitation is that the NCH data are from a database of clinical encounters. Based on comparison to CHAT, all primary age-related changes appeared qualitatively (and often quantitatively) conserved across studies, suggesting that these robust effects reflect fundamental developmental processes that may transcend diagnostic status. The subjects in CHAT either qualified for a diagnosis of obstructive sleep apnea, or snored and on that basis had some form of sleep-disordered breathing, meaning that the subjects though not necessarily referred to a sleep disorders center still most likely did not have normal sleep. Therefore, our comparisons may be limited due to lack of truly normative controls. Another limitation is the absence of detailed cognitive and behavioral data in the clinical cohort, precluding more direct investigations of how age-related changes in sleep track with development. Similarly, puberty status was not available, limiting our ability to interpret developmental differences not captured by age. Nonetheless, the presence of sleep-disordered breathing or other sleep disorders in many of the subjects who contributed data for the present analyses does not invalidate the high likelihood that sleep in these individuals still reflected many aspects of normal sleep development across childhood and adolescence. For example, we did not find evidence of significant effects of AHI on the age-related changes of sleep metrics or brain age prediction.

Our findings from retrospective clinical and clinical research data, meanwhile, appear to suggest very strong age-related changes across numerous sleep metrics in children ages 4 to 17 years, which could be robustly identified across independent samples despite the demographic, clinical and procedural differences. As well as confirming previous reports based on smaller samples, we describe new metrics not previously studied from a developmental perspective, including stage- and channel-specific developmental trajectories of relative spectral power, intra-spindle frequency modulation, and temporal overlap between spindles and SOs. A model of multiple sleep metrics across different domains was able to predict chronological age with high precision in typically developing individuals, whereas this correspondence was lessened in individuals with NDDs, suggesting that these sleep metrics are sensitive to various functional abnormalities present in the (sleeping) brain. Taken together, our results indicated that sleep macro and microarchitecture offer important information about brain maturation which may facilitate a better understanding of the atypical neurodevelopment.

## METHODS

### Participants

We primarily used PSG data from two pediatric samples – the Nationwide Children’s Hospital Sleep DataBank (NCH) and the Child Adenotonsillectomy Trial (CHAT) – both available via the National Sleep Research Resource (http://sleepdata.org). The NCH sample was composed of patients (from infants to some adults) who underwent clinical PSG (Lee et al., 2022), and contained diagnostic (ICD 9/10 codes) and medication data. All the data were de-identified prior to NSRR deposition, and received NCH Institutional Review Board exemption with HIPAA waiver.

The CHAT sample was derived from data collected from six US pediatric clinical centers as part of sleep screening procedures for a clinical trial of children aged 5 to 9 with snoring who were candidates for adenotonsillectomy. All participants were without severe chronic medical conditions or ADHD requiring medications, presented with snoring and were potential candidates for adenotonsillectomy (Marcus et al., 2013; Weinstock et al., 2014). All children (n=1,244) were screened with a PSG, with 464 who met the eligibility criteria (AHI 2-30 and presence of mild obstructive sleep apnea (OSA), for detailed description of inclusion criteria see Marcus et. al., 2013) being enrolled in the study. Here we used the sample of all screened children including non-randomized sample without OSA (AHI < 3) and those with severe OSA (AHI > 30), but excluded the follow-up recordings that had been performed on participants with mild to moderate obstructive apnea. Data collection for CHAT was approved by local Institutional Review Boards and written informed consent was obtained from each individual or their legal guardians.

For the brain age analysis analyses only, to expand the age range of the test set, we additionally included individuals from the Pediatric Adenotonsillectomy Trial for Snoring (PATS) dataset. This allowed us to test the transferability of the model (Wang et al., 2020). For the PATS study, children with symptoms of obstructive sleep-disordered breathing were included. The study was similar to CHAT, but included a wider age range, and specifically focused on children with snoring but with an AHI <3.

Cleveland Family Study sample (including individuals between 6 and 88 years) was used for analysis testing the possibility of accelerated aging in the DS subgroup.

### Clinical information for NCH sample

The DIAGNOSIS.csv file (available via NSRR) was used to delineate clinical sub-groups in the NCH sample. Following recommendations from the original description of the dataset (Lee et al., 2021), we only used final diagnosis codes (DX_ENC_TYPE & DX_SOURCE_TYPE columns equal to “Final Dx”). Since diagnostic codes provided for the sample were either according ICD9 or ICD10, we searched for specific diagnoses using the string search based on the diagnosis description (DX_NAME). For example, searching a string “[Aa]utis” and visually checking all unique matching diagnoses as well as the ICD codes. The information for all matching diagnoses for each condition is provided in the Supplementary table 1.

To control for medication use, we used records available in the MEDICATION.csv file. Specifically, we identified participants whose PSG was performed between the prescribed medication start and end date. We identified four therapeutic classes of medication out of 42 that could potentially affect sleep: 1) antihistamines, 2) psychotherapeutic drugs, 3) CNS drugs, including anticonvulsants, 4) hormones, and 5) sedatives/hypnotics. We summarized them by therapeutic class and subclass (THERA_CLASS and THERA_SUBCLASS), pharmaceutical class (PHARMA_CLASS).

### EEG preprocessing

All steps of sleep EEG data processing were performed using Luna (http://zzz.bwh.harvard.edu/luna/), an open-source package developed by us (S.M.P). All NCH, CHAT and PATS PSGs contained six EEG channels (F3, F4, C3, C4, O1, O2). In CHAT, two temporal channels (T3, T4) were also available. The CFS cohort contained C3 and C4 EEG channels only. We first selected 30-seccond epochs of a particular stage (N2, N3, REM) according to manual, AASM-based staging. All EEG signals were down-sampled to 200 Hz, referenced to contralateral mastoids (M1 or M2), converted to uV units and bandpass filtered between 0.5 and 35 Hz. Due to excessive line noise interference observed in many NCH samples, we applied an approach to remove it based on spectrum interpolation (Leske & Dalal, 2019), as implemented in Luna.

Next, within each stage, we identified all epochs with maximum amplitudes above 200 uV, or with flat or clipped signals for more than 10% of the epoch; further, epochs were marked as outliers if they were i) more than 3 SDs from the mean (for that individual) of all channels for any of the three Hjorth parameters, activity, mobility and complexity, ii) 4 SDs from the mean of other epochs of the same channel or iii) 4 SDs from the mean of all epochs across all channels. Hjorth-based epoch outlier removal was performed twice for each individual. Channels and/or epochs were removed if more than 50% of epochs were outliers.

### Spectral power estimation

Spectral power was estimated using Welch’s method separately for N2, N3 and REM, summarized by classical frequency bands – slow (0.5-1 Hz), delta (1-4 Hz), theta (4-8 Hz), alpha (8-12 Hz), sigma (12-15 Hz), beta (15-30 Hz), and total power (0.5 to 50 Hz). Specifically, for each 30-seccond epoch, we applied the Fast Fourier Transform with 4 second segments (0.25 Hz spectral resolution) windowing with a Tukey (50%) taper, with consecutive segments overlapping by 50% (2 seconds). We then averaged power across all segments per epoch. Subsequently, epoch-wise power was averaged across all epochs for a particular channel and stage. Relative power was computed with respect to the total absolute power. Absolute power values were then log-transformed prior to analysis.

### Spindle detection

Motivated by recent findings that two classes of spindles – slow frontal and fast central – emerge as early as 18 months after birth (Kwon et al., 2022), we detected both separately. Slow and fast spindles were detected using center frequencies of 11Hz and 15Hz respectively using a wavelet method as previously described (Purcell et al., 2017). Specifically, putative spindles were identified based on temporally smoothed (window duration=0.1 s) wavelet coefficients (from a complex Morlet wavelet transform) using following criteria. Intervals exceeding 1) 4.5 times the mean for at least 300 ms and also 2) 2 times the mean for at least 500ms were selected as putative spindles. Intervals over 3 seconds were rejected; consecutive intervals within 500ms were merged (unless the resulting spindle was greater than 3 seconds). In addition, putative spindles were discarded if the relative increase in non-spindle bands activity (delta, theta, and beta) was greater than the relative increase in spindle frequency activity (i.e. relative to all N2 sleep), thereby ensuring putative spindles preferentially reflect sigma band activity, and not general increases in signal amplitude, which is often due to artifact or other non-spindle activities. Based on the set of spindles that passed QC, we computed spindle density (count per minute), amplitude, duration, observed frequency, and chirp (intra-spindle frequency change).

### SO detection

Zero-crossings were identified based on the EEG signals band-pass filtered between 0.5 and 4 Hz. To define putative SO the following temporal criteria were satisfied: 1) a consecutive zero-crossing leading to negative peak was between 0.3-1.5 seconds; 2) a zero-crossing leading to positive peak were not longer than 1 second. With respect to amplitude criteria, two separate approaches were used, similar to Djonlagic et al (2020). First, an adaptive/relative threshold (our default) such that negative peak and peak-to-peak amplitudes were required to be greater than twice the mean (for that individual/channel). Second, an absolute threshold requiring a negative peak amplitude larger -40 uV, and peak-to-peak amplitude larger then 75 uV. SO density (count per minute) as well as the mean amplitude of the negative peak, peak-to-peak amplitude, duration and the upward slope of negative peak were estimated for each channel.

### Coupling between SO and spindles

For each channel we identified spindles that overlapped with detected SO and characterized their coupling using the following three metrics. First, we computed the proportion of spindles that overlapped with a SO (“gross overlap”). Using the filter-Hilbert method, we also estimated SO phase at the spindle peak, which was averaged (circular mean) across SOs for each channel (coupling angle at spindle peak). In addition, the inter-trial phase clustering assessed the consistency of non-uniform phase coupling between SO and spindles (coupling magnitude). Overlap and magnitude metrics were z-transformed using a null distribution of same metrics generated during 10,000 random permutations where time indices of the time series were shuffled in a manner that preserved the overall number of SOs, spindles and the gross overlap between SO and spindles (the latter is true only for the coupling magnitude).

### Exclusion criteria

Primary exclusion criteria for the NCH sample were i) age younger than 2.5 years (due to potential differences in infant and toddler EEGs), ii) age above 17.5 years (due to sparsity of data, and iii) a narcolepsy diagnosis (n=42). In CHAT, we removed individuals for whom age information was missing (N=19). If the same individual had multiple recordings available, we only used a single (the first) recording.

An additional exclusion criterion applied for the macro-architecture analysis was TST < 180 mins. For analysis of spectral power and spindles, additional exclusion criteria were applied: i) N of available epochs for each stage (N2, N3, REM) after outlier removal less than 10, ii) persistent line noise interference (SPK measure > 5 SD in least one channel at any stage), iii) outlier spectral power at 1 Hz (< 4 SD or > 4 SD in least one channel at any stage) to target movement, ocular artifacts, general low signal to noise ratio, or at 25 Hz (> 4 SD in least one channel at any stage) to target muscle activity artifacts.

Signal polarity flips were observed in a portion of recordings in both samples (for details on polarity in several NSRR samples: https://zzz.bwh.harvard.edu/luna/vignettes/nsrr-polarity/) and additional exclusion criteria were necessary for analyses dependent on signal polarity – those involving SO and coupling between SO and spindles. Recordings with ambiguous polarity were removed for these analyses (−1 < T_DIFF < 1 at C3 or C4 during N2 stage from Luna’s POL command) and polarity of all recordings with T_DIFF > 1 at C3 or C4 during N2 stage was flipped.

Final sample size and demographic characteristics for the primary samples used (for NCH and CHAT) are in Table 1. The same exclusion criteria were applied to PATS. Characteristics of the final analytic sample that was included for brain age prediction were: total N = 587, with 285 females, 354 white, 189 black or African American individuals, with a mean age of 6.3 years (range 3 – 12.8 years).

### Statistical analysis

We used a linear regression models of each sleep metric regressed on age, also controlling for sex and race/ethnicity. Outliers (using a 3 SD criterion) were removed for each sleep metric (repeated twice). In a control analysis we added Apnea-hypopnea index (AHI) computed as an average number of apnea and/or hypopnea events per hour of sleep as an additional covariate to linear regression model, which yielded almost identical results (Pearson correlation between signed -log 10 p-value controlling and not controlling for AHI was > 0.99 in both NCH and CHAT samples). To describe effect sizes of age-related changes in sleep metrics, we also calculated Pearson correlations between sleep metrics and age.

In analyses of NCH clinical subgroups, we used linear regression model controlling for race, sex, AHI, co-occurring sleep diagnoses, other NDD diagnoses and medication use. For **Sup. Figure 5**, prior to running a linear regression, 3-SD outliers were removed in two rounds and sleep metrics’ estimates were z-transformed to obtained standardized linear regression coefficients. P-values were adjusted for multiple comparisons (all sleep estimates n=321) separately for each subgroup and tested effect (subgroup, subgroup by age interaction) using FDR method (Benjamini & Yekutieli, 2001).

To predict individuals’ ages, we trained a multiple linear regression model using the sleep metrics studied here. After excluding subjects from clinical subgroups, we randomly split the NCH sample into training (70% of subjects) and testing (30%) sets. CHAT, PATS and NCH clinical subgroups sample were retained as additional, independent testing sets. To reduce the number of features for prediction model, we only kept those that were significantly (p<10^−10^) associated with age in the training dataset. In addition, for features that were highly correlated (abs r > 0.9), one was excluded. Remaining features (for all datasets) were z-transformed using the mean and standard deviation of the training set. Sex and race were included as covariates in all models.

We additionally applied an alternative to conventional linear regression: specifically, gradient descent boosting machines as implemented in the LightGBM machine learning library. Performance was near identical to the linear regression model, in this particular case, and so we retained the simpler model, as performance in terms of age prediction was already high (data not shown).

To estimate the variability in the prediction of results, we repeated the step of fitting the model 50 times with different, non-overlapping individuals from NCH randomly assigned to testing and training set each round. The other testing sets (CHAT, PATS and clinical subgroups) were kept the same. Estimates of model performance – Pearson’s correlation, mean absolute error and mean error between predicted and true age – were then averaged across 50 rounds and min and max values were estimated. We used a two-sample t-test to test if there was significant difference in MAE and ME between NCH testing set and each clinical subgroup in each round and reported the median p-value in **Figure 4 B**.

We applied the following steps to estimate similarity in all sleep variables (macroarchitecture, absolute and relative band power, spindles, SO estimates and coupling metrics) across different age bins of combined sample of PATS and CFS cohorts and DS group (**Figure 5H)**. First, we z-scored all sleep variables across individuals of the combined sample and NCH sample using mean and standard deviation of the combined sample of each sleep metric Then, we defined age bins in the combined sample (two-year non-overlapping windows centered at 3, 5, 7, 9, 11, 13 and four-year non-overlapping windows centered at 16, 20, …, 80 years). Then means were computed for each sleep metric across individuals belonging to a particular age bin in the combined sample, as well as the DS subgroup and non-NDD group of the NCH sample. Finally, the average absolute difference between DS (and non-NDD) means and each age bin means were computed and plotted as a function of age.

## REFERENCES

Andrillon, T., Nir, Y., Staba, R. J., Ferrarelli, F., Cirelli, C., Tononi, G., & Fried, I. (2011). Sleep Spindles in Humans: Insights from Intracranial EEG and Unit Recordings. The Journal of Neuroscience, 31(49), 17821–17834. https://doi.org/10.1523/JNEUROSCI.2604-11.2011

Baecker, L., Garcia-Dias, R., Vieira, S., Scarpazza, C., & Mechelli, A. (2021). Machine learning for brain age prediction: Introduction to methods and clinical applications. EBioMedicine, 72, 103600. https://doi.org/10.1016/j.ebiom.2021.103600

Baker, F. C., Turlington, S. R., & Colrain, I. (2012). Developmental changes in the sleep electroencephalogram of adolescent boys and girls. Journal of Sleep Research, 21(1), 59–67. https://doi.org/10.1111/j.1365-2869.2011.00930.x

Baker, F. C., Willoughby, A. R., Massimiliano, de Z., Franzen, P. L., Prouty, D., Javitz, H., Hasler, B., Clark, D. B., & Colrain, I. M. (2016). Age-Related Differences in Sleep Architecture and Electroencephalogram in Adolescents in the National Consortium on Alcohol and Neurodevelopment in Adolescence Sample. Sleep, 39(7), 1429–1439. https://doi.org/10.5665/sleep.5978

Baker, F. C., Yűksel, D., & de Zambotti, M. (2020). Sex Differences in Sleep. In H. Attarian & M. Viola-Saltzman (Eds.), Sleep Disorders in Women: A Guide to Practical Management (pp. 55–64). Springer International Publishing. https://doi.org/10.1007/978-3-030-40842-8_5

Barriga-Paulino, C. I., Flores, A. B., & Gómez, C. M. (2011). Developmental changes in the EEG rhythms of children and young adults: Analyzed by means of correlational, brain topography and principal component analysis. Journal of Psychophysiology, 25(3), 143–158. https://doi.org/10.1027/0269-8803/a000052

Becker, S. P. (2020). ADHD and sleep: Recent advances and future directions. Current Opinion in Psychology, 34, 50–56. https://doi.org/10.1016/j.copsyc.2019.09.006

Benjamini, Y., & Yekutieli, D. (2001). The control of the false discovery rate in multiple testing under dependency. The Annals of Statistics, 29(4), 1165–1188. https://doi.org/10.1214/aos/1013699998

Bocskai, G., Pótári, A., Gombos, F., & Kovács, I. (2022). The adolescent pattern of sleep spindle development revealed by HD-EEG. Journal of Sleep Research, n/a(n/a), e13618. https://doi.org/10.1111/jsr.13618

Boord, P. R., Rennie, C. J., & Williams, L. M. (2007). INTEGRATING “BRAIN” AND “BODY” MEASURES: CORRELATIONS BETWEEN EEG AND METABOLIC CHANGES OVER THE HUMAN LIFESPAN. Journal of Integrative Neuroscience, 06(01), 205–218. https://doi.org/10.1142/S0219635207001416

Buchmann, A., Ringli, M., Kurth, S., Schaerer, M., Geiger, A., Jenni, O. G., & Huber, R. (2011). EEG Sleep Slow-Wave Activity as a Mirror of Cortical Maturation. Cerebral Cortex, 21(3), 607–615. https://doi.org/10.1093/cercor/bhq129

Campbell, I. G., & Feinberg, I. (2016). Maturational Patterns of Sigma Frequency Power Across Childhood and Adolescence: A Longitudinal Study. Sleep, 39(1), 193–201. https://doi.org/10.5665/sleep.5346

Campbell, I. G., Grimm, K. J., de Bie, E., & Feinberg, I. (2012). Sex, puberty, and the timing of sleep EEG measured adolescent brain maturation. Proceedings of the National Academy of Sciences of the United States of America, 109(15), 5740– 5743. https://doi.org/10.1073/pnas.1120860109

Carvalho, D. Z., Gerhardt, G. J. L., Dellagustin, G., de Santa-Helena, E. L., Lemke, N., Segal, A. Z., & Schönwald, S. V. (2014). Loss of sleep spindle frequency deceleration in Obstructive Sleep Apnea. Clinical Neurophysiology, 125(2), 306– 312. https://doi.org/10.1016/j.clinph.2013.07.005

Cole, J. H., Annus, T., Wilson, L. R., Remtulla, R., Hong, Y. T., Fryer, T. D., Acosta-Cabronero, J., Cardenas-Blanco, A., Smith, R., Menon, D. K., Zaman, S. H., Nestor, P. J., & Holland, A. J. (2017). Brain-predicted age in Down syndrome is associated with beta amyloid deposition and cognitive decline. Neurobiology of Aging, 56, 41–49. https://doi.org/10.1016/j.neurobiolaging.2017.04.006

Cole, J. H., Leech, R., Sharp, D. J., & Initiative, for the A. D. N. (2015). Prediction of brain age suggests accelerated atrophy after traumatic brain injury. Annals of Neurology, 77(4), 571–581. https://doi.org/10.1002/ana.24367

Dajani, D. R., Llabre, M. M., Nebel, M. B., Mostofsky, S. H., & Uddin, L. Q. (2016). Heterogeneity of executive functions among comorbid neurodevelopmental disorders. Scientific Reports, 6(1), Article 1. https://doi.org/10.1038/srep36566

Djonlagic, I., Mariani, S., Fitzpatrick, A. L., Van Der Klei, V. M. G. T. H., Johnson, D. A., Wood, A. C., Seeman, T., Nguyen, H. T., Prerau, M. J., Luchsinger, J. A., Dzierzewski, J. M., Rapp, S. R., Tranah, G. J., Yaffe, K., Burdick, K. E., Stone, K. L., Redline, S., & Purcell, S. M. (2020). Macro and micro sleep architecture and cognitive performance in older adults. Nature Human Behaviour. https://doi.org/10.1038/s41562-020-00964-y

Dustman, R. E., Shearer, D. E., & Emmerson, R. Y. (1999). Life-span changes in EEG spectral amplitude, amplitude variability and mean frequency. Clinical Neurophysiology, 110(8), 1399–1409. https://doi.org/10.1016/S1388-2457(99)00102-9

Esposito, M., & Carotenuto, M. (2014). Intellectual disabilities and power spectra analysis during sleep: A new perspective on borderline intellectual functioning. Journal of Intellectual Disability Research, 58(5), 421–429. https://doi.org/10.1111/jir.12036

Feinberg, I., & Campbell, I. G. (2013). Longitudinal sleep EEG trajectories indicate complex patterns of adolescent brain maturation. American Journal of Physiology-Regulatory, Integrative and Comparative Physiology, 304(4), R296–R303. https://doi.org/10.1152/ajpregu.00422.2012

Feinberg, I., Davis, N. M., de Bie, E., Grimm, K. J., & Campbell, I. G. (2012). The maturational trajectories of NREM and REM sleep durations differ across adolescence on both school-night and extended sleep. American Journal of Physiology-Regulatory, Integrative and Comparative Physiology, 302(5), R533–R540. https://doi.org/10.1152/ajpregu.00532.2011

Fernandez, L. M. J., & Lüthi, A. (2020). Sleep Spindles: Mechanisms and Functions. Physiological Reviews, 100(2), 805–868. https://doi.org/10.1152/physrev.00042.2018

Franke, K., & Gaser, C. (2019). Ten Years of BrainAGE as a Neuroimaging Biomarker of Brain Aging: What Insights Have We Gained? Frontiers in Neurology, 10. https://www.frontiersin.org/articles/10.3389/fneur.2019.00789

Franke, K., Luders, E., May, A., Wilke, M., & Gaser, C. (2012). Brain maturation: Predicting individual BrainAGE in children and adolescents using structural MRI. NeuroImage, 63(3), 1305–1312. https://doi.org/10.1016/j.neuroimage.2012.08.001

Galland, B. C., Taylor, B. J., Elder, D. E., & Herbison, P. (2012). Normal sleep patterns in infants and children: A systematic review of observational studies. Sleep Medicine Reviews, 16(3), 213–222. https://doi.org/10.1016/j.smrv.2011.06.001

Gameren-Oosterom, H. B. M. van, Fekkes, M., Buitendijk, S. E., Mohangoo, A. D., Bruil, J., & Wouwe, J. P. V. (2011). Development, Problem Behavior, and Quality of Life in a Population Based Sample of Eight-Year-Old Children with Down Syndrome. PLOS ONE, 6(7), e21879. https://doi.org/10.1371/journal.pone.0021879

Gandal, M. J., Leppa, V., Won, H., Parikshak, N. N., & Geschwind, D. H. (2016). The road to precision psychiatry: Translating genetics into disease mechanisms. Nature Neuroscience, 19(11), 1397–1407. https://doi.org/10.1038/nn.4409

Gasser, T., Verleger, R., Bächer, P., & Sroka, L. (1988). Development of the EEG of school-age children and adolescents. I. Analysis of band power. Electroencephalography and Clinical Neurophysiology, 69(2), 91–99. https://doi.org/10.1016/0013-4694(88)90204-0

Gaudreau, H., Carrier, J., & Montplaisir, J. (2001). Age-related modifications of NREM sleep EEG: From childhood to middle age. Journal of Sleep Research, 10(3), 165–172. https://doi.org/10.1046/j.1365-2869.2001.00252.x

Goldstone, A., Willoughby, A. R., de Zambotti, M., Clark, D. B., Sullivan, E. V., Hasler, B. P., Franzen, P. L., Prouty, D. E., Colrain, I. M., & Baker, F. C. (2019). Sleep spindle characteristics in adolescents. Clinical Neurophysiology, 130(6), 893– 902. https://doi.org/10.1016/j.clinph.2019.02.019

Goldstone, A., Willoughby, A. R., de Zambotti, M., Franzen, P. L., Kwon, D., Pohl, K. M., Pfefferbaum, A., Sullivan, E. V., Müller-Oehring, E. M., Prouty, D. E., Hasler, B. P., Clark, D. B., Colrain, I. M., & Baker, F. C. (2018). The mediating role of cortical thickness and gray matter volume on sleep slow-wave activity during adolescence. Brain Structure and Function, 223(2), 669–685. https://doi.org/10.1007/s00429-017-1509-9

Gorgoni, M., Scarpelli, S., Reda, F., & De Gennaro, L. (2020). Sleep EEG oscillations in neurodevelopmental disorders without intellectual disabilities. Sleep Medicine Reviews, 49, 101224. https://doi.org/10.1016/j.smrv.2019.101224

Hahn, M. A., Heib, D., Schabus, M., Hoedlmoser, K., & Helfrich, R. F. (2020). Slow oscillation-spindle coupling predicts enhanced memory formation from childhood to adolescence. ELife, 9, e53730. https://doi.org/10.7554/eLife.53730

Hayashi, M., Inoue, Y., Iwakawa, Y., & Sasaki, H. (1990). REM sleep abnormalities in severe athetoid cerebral palsy. Brain and Development, 12(5), 494–497. https://doi.org/10.1016/S0387-7604(12)80214-2

Hoedlmoser, K., Heib, D. P. J., Roell, J., Peigneux, P., Sadeh, A., Gruber, G., & Schabus, M. (2014). Slow Sleep Spindle Activity, Declarative Memory, and General Cognitive Abilities in Children. Sleep, 37(9), 1501–1512. https://doi.org/10.5665/sleep.4000

Hong, J., Feng, Z., Wang, S.-H., Peet, A., Zhang, Y.-D., Sun, Y., & Yang, M. (2020). Brain Age Prediction of Children Using Routine Brain MR Images via Deep Learning. Frontiers in Neurology, 11. https://www.frontiersin.org/articles/10.3389/fneur.2020.584682

Horvath, S., Garagnani, P., Bacalini, M. G., Pirazzini, C., Salvioli, S., Gentilini, D., Di Blasio, A. M., Giuliani, C., Tung, S., Vinters, H. V., & Franceschi, C. (2015). Accelerated epigenetic aging in Down syndrome. Aging Cell, 14(3), 491–495. https://doi.org/10.1111/acel.12325

Iglowstein, I., Jenni, O. G., Molinari, L., & Largo, R. H. (2003). Sleep Duration From Infancy to Adolescence: Reference Values and Generational Trends. Pediatrics, 111(2), 302–307. https://doi.org/10.1542/peds.111.2.302

Jenni, O. G., & Carskadon, M. A. (2004). Spectral Analysis of the Sleep Electroencephalogram During Adolescence. Sleep, 27(4), 774–783. https://doi.org/10.1093/sleep/27.4.774

Jeste, S. S., Frohlich, J., & Loo, S. K. (2015). Electrophysiological biomarkers of diagnosis and outcome in neurodevelopmental disorders. Current Opinion in Neurology, 28(2), 110–116. https://doi.org/10.1097/WCO.0000000000000181

Kamara, D., & Beauchaine, T. P. (2020). A Review of Sleep Disturbances among Infants and Children with Neurodevelopmental Disorders. Review Journal of Autism and Developmental Disorders, 7(3), 278–294. https://doi.org/10.1007/s40489-019-00193-8

Knoop, M. S., de Groot, E. R., & Dudink, J. (2021). Current ideas about the roles of rapid eye movement and non–rapid eye movement sleep in brain development. Acta Paediatrica, 110(1), 36–44. https://doi.org/10.1111/apa.15485

Kopasz, M., Loessl, B., Hornyak, M., Riemann, D., Nissen, C., Piosczyk, H., & Voderholzer, U. (2010). Sleep and memory in healthy children and adolescents – A critical review. Sleep Medicine Reviews, 14(3), 167–177. https://doi.org/10.1016/j.smrv.2009.10.006

Kurth, S., Jenni, O. G., Riedner, B. A., Tononi, G., Carskadon, M. A., & Huber, R. (2010). Characteristics of Sleep Slow Waves in Children and Adolescents. Sleep, 33(4), 475–480. https://doi.org/10.1093/sleep/33.4.475

Kurth, S., Ringli, M., Geiger, A., LeBourgeois, M., Jenni, O. G., & Huber, R. (2010). Mapping of Cortical Activity in the First Two Decades of Life: A High-Density Sleep Electroencephalogram Study. Journal of Neuroscience, 30(40), 13211– 13219. https://doi.org/10.1523/JNEUROSCI.2532-10.2010

Kwon, H., Walsh, K. G., Berja, E. D., Manoach, D. S., Eden, U. T., Kramer, M. A., & Chu, C. J. (2022). Normative sleep spindle database and findings from 772 healthy children from birth through 18 years (p. 2022.03.31.486476). bioRxiv. https://doi.org/10.1101/2022.03.31.486476

Larson, A. M., Ryther, R. C. C., Jennesson, M., Geffrey, A. L., Bruno, P. L., Anagnos, C. J., Shoeb, A. H., Thibert, R. L., & Thiele, E. A. (2012). Impact of pediatric epilepsy on sleep patterns and behaviors in children and parents. Epilepsia, 53(7), 1162– 1169. https://doi.org/10.1111/j.1528-1167.2012.03515.x

Leary, E. B., Watson, K. T., Ancoli-Israel, S., Redline, S., Yaffe, K., Ravelo, L. A., Peppard, P. E., Zou, J., Goodman, S. N., Mignot, E., & Stone, K. L. (2020). Association of Rapid Eye Movement Sleep With Mortality in Middle-aged and Older Adults. JAMA Neurology, 77(10), 1241–1251. https://doi.org/10.1001/jamaneurol.2020.2108

Lee, H., Li, B., DeForte, S., Splaingard, M., Huang, Y., Chi, Y., & Linwood, S. L. (2021). NCH Sleep DataBank: A Large Collection of Real-world Pediatric Sleep Studies. ArXiv:2102.13284 [Eess, Stat]. http://arxiv.org/abs/2102.13284

Lee, H., Li, B., DeForte, S., Splaingard, M. L., Huang, Y., Chi, Y., & Linwood, S. L. (2022). A large collection of real-world pediatric sleep studies. Scientific Data, 9(1), Article 1. https://doi.org/10.1038/s41597-022-01545-6

Leske, S., & Dalal, S. S. (2019). Reducing power line noise in EEG and MEG data via spectrum interpolation. NeuroImage, 189, 763–776. https://doi.org/10.1016/j.neuroimage.2019.01.026

Li, H., Satterthwaite, T. D., & Fan, Y. (2018). Brain age prediction based on resting-state functional connectivity patterns using convolutional neural networks. 2018 IEEE 15th International Symposium on Biomedical Imaging (ISBI 2018), 101–104. https://doi.org/10.1109/ISBI.2018.8363532

Lund, M. J., Alnæs, D., de Lange, A.-M. G., Andreassen, O. A., Westlye, L. T., & Kaufmann, T. (2022). Brain age prediction using fMRI network coupling in youths and associations with psychiatric symptoms. NeuroImage: Clinical, 33, 102921. https://doi.org/10.1016/j.nicl.2021.102921

Marcus, C. L., Moore, R. H., Rosen, C. L., Giordani, B., Garetz, S. L., Taylor, H. G., Mitchell, R. B., Amin, R., Katz, E. S., Arens, R., Paruthi, S., Muzumdar, H., Gozal, D., Thomas, N. H., Ware, J., Beebe, D., Snyder, K., Elden, L., Sprecher, R. C., … Childhood Adenotonsillectomy Trial (CHAT). (2013). A randomized trial of adenotonsillectomy for childhood sleep apnea. The New England Journal of Medicine, 368(25), 2366–2376. https://doi.org/10.1056/NEJMoa1215881

Nakamura, E., & Tanaka, S. (1998). Biological ages of adult men and women with Down’s syndrome and its changes with aging. Mechanisms of Ageing and Development, 105(1), 89–103. https://doi.org/10.1016/S0047-6374(98)00081-5

Nygate, Y., Rusk, S., Fernandez, C., Glattard, N., Arguelles, J., Shi, J., Hwang, D., & Watson, N. (2021). 543 EEG-Based Deep Neural Network Model for Brain Age Prediction and Its Association with Patient Health Conditions. Sleep, 44(Supplement_2), A214. https://doi.org/10.1093/sleep/zsab072.541

Ohayon, M. M., Carskadon, M. A., Guilleminault, C., & Vitiello, M. V. (2004). Meta-Analysis of Quantitative Sleep Parameters From Childhood to Old Age in Healthy Individuals: Developing Normative Sleep Values Across the Human Lifespan. Sleep, 27(7), 1255–1273. https://doi.org/10.1093/sleep/27.7.1255

Pack, A. M. (2019). Epilepsy Overview and Revised Classification of Seizures and Epilepsies. CONTINUUM: Lifelong Learning in Neurology, 25(2), 306. https://doi.org/10.1212/CON.0000000000000707

Purcell, S. M., Manoach, D. S., Demanuele, C., Cade, B. E., Mariani, S., Cox, R., Panagiotaropoulou, G., Saxena, R., Pan, J. Q., Smoller, J. W., Redline, S., & Stickgold, R. (2017). Characterizing sleep spindles in 11,630 individuals from the National Sleep Research Resource. Nature Communications, 8(1), Article 1. https://doi.org/10.1038/ncomms15930

Qin, J., Chen, S.-G., Hu, D., Zeng, L.-L., Fan, Y.-M., Chen, X.-P., & Shen, H. (2015). Predicting individual brain maturity using dynamic functional connectivity. Frontiers in Human Neuroscience, 9. https://www.frontiersin.org/article/10.3389/fnhum.2015.00418

Robinson-Shelton, A., & Malow, B. A. (2015). Sleep Disturbances in Neurodevelopmental Disorders. Current Psychiatry Reports, 18(1), 6. https://doi.org/10.1007/s11920-015-0638-1

Rosello, M., Caro-Llopis, A., Orellana, C., Oltra, S., Alemany-Albert, M., Marco-Hernandez, A. V., Monfort, S., Pedrola, L., Martinez, F., & Tomás, M. (2021). Hidden etiology of cerebral palsy: Genetic and clinical heterogeneity and efficient diagnosis by next-generation sequencing. Pediatric Research, 90(2), Article 2. https://doi.org/10.1038/s41390-020-01250-3

Sadak, U., Honrath, P., Ermis, U., Heckelmann, J., Meyer, T., Weber, Y., & Wolking, S. (2022). Reduced REM sleep: A potential biomarker for epilepsy – a retrospective case-control study. Seizure, 98, 27–33. https://doi.org/10.1016/j.seizure.2022.03.022

Scholle, S., Beyer, U., Bernhard, M., Eichholz, S., Erler, T., Graneß, P., Goldmann-Schnalke, B., Heisch, K., Kirchhoff, F., Klementz, K., Koch, G., Kramer, A., Schmidtlein, C., Schneider, B., Walther, B., Wiater, A., & Scholle, H. C. (2011). Normative values of polysomnographic parameters in childhood and adolescence: Quantitative sleep parameters. Sleep Medicine, 12(6), 542–549. https://doi.org/10.1016/j.sleep.2010.11.011

Scholle, S., Zwacka, G., & Scholle, H. C. (2007). Sleep spindle evolution from infancy to adolescence. Clinical Neurophysiology, 118(7), 1525–1531. https://doi.org/10.1016/j.clinph.2007.03.007

Segalowitz, S. J., Santesso, D. L., & Jetha, M. K. (2010). Electrophysiological changes during adolescence: A review. Brain and Cognition, 72(1), 86–100. https://doi.org/10.1016/j.bandc.2009.10.003

Shibagaki, M., & Kiyono, S. (1983). Duration of spindle bursts during nocturnal sleep in mentally retarded children. Electroencephalography and Clinical Neurophysiology, 55(6), 645–651. https://doi.org/10.1016/0013-4694(83)90274-2

Shibagaki, M., Kiyono, S., & Watanabe, K. (1980). Nocturnal sleep in severely mentally retarded children: Abnormal eeg patterns in sleep cycle. Electroencephalography and Clinical Neurophysiology, 49(3), 337–344. https://doi.org/10.1016/0013-4694(80)90228-X

Shinomiya, S., Nagata, K., Takahashi, K., & Masumura, T. (1999). Development of Sleep Spindles in Young Children and Adolescents. Clinical Electroencephalography, 30(2), 39–43. https://doi.org/10.1177/155005949903000203

Sibarani, C. R., Walter, L. M., Davey, M. J., Nixon, G. M., & Horne, R. S. C. (2022). Sleep-disordered breathing and sleep macro- and micro-architecture in children with Down syndrome. Pediatric Research, 91(5), Article 5. https://doi.org/10.1038/s41390-021-01642-z

Simard-Tremblay, E., Constantin, E., Gruber, R., Brouillette, R. T., & Shevell, M. (2011). Sleep in Children With Cerebral Palsy: A Review. Journal of Child Neurology, 26(10), 1303–1310. https://doi.org/10.1177/0883073811408902

Soroko, S. I., Shemyakina, N. V., Nagornova, Zh. V., & Bekshaev, S. S. (2014). Longitudinal study of EEG frequency maturation and power changes in children on the Russian North. International Journal of Developmental Neuroscience, 38, 127–137. https://doi.org/10.1016/j.ijdevneu.2014.08.012

Souders, M. C., Zavodny, S., Eriksen, W., Sinko, R., Connell, J., Kerns, C., Schaaf, R., & Pinto-Martin, J. (2017). Sleep in Children with Autism Spectrum Disorder. Current Psychiatry Reports, 19(6), 34. https://doi.org/10.1007/s11920-017-0782-x

Spanò, G., Gómez, R. L., Demara, B. I., Alt, M., Cowen, S. L., & Edgin, J. O. (2018). REM sleep in naps differentially relates to memory consolidation in typical preschoolers and children with Down syndrome. Proceedings of the National Academy of Sciences, 115(46), 11844–11849. https://doi.org/10.1073/pnas.1811488115

Stiles, J., & Jernigan, T. L. (2010). The Basics of Brain Development. Neuropsychology Review, 20(4), 327–348. https://doi.org/10.1007/s11065-010-9148-4

Stores, G., & Stores, R. (2013). Sleep disorders and their clinical significance in children with Down syndrome. Developmental Medicine & Child Neurology, 55(2), 126– 130. https://doi.org/10.1111/j.1469-8749.2012.04422.x

Sun, H., Paixao, L., Oliva, J. T., Goparaju, B., Carvalho, D. Z., van Leeuwen, K. G., Akeju, O., Thomas, R. J., Cash, S. S., Bianchi, M. T., & Westover, M. B. (2019). Brain age from the electroencephalogram of sleep. Neurobiology of Aging, 74, 112–120. https://doi.org/10.1016/j.neurobiolaging.2018.10.016

Surtees, A. D. R., Oliver, C., Jones, C. A., Evans, D. L., & Richards, C. (2018). Sleep duration and sleep quality in people with and without intellectual disability: A meta-analysis. Sleep Medicine Reviews, 40, 135–150. https://doi.org/10.1016/j.smrv.2017.11.003

Tarokh, L., & Carskadon, M. A. (2010). Developmental Changes in the Human Sleep EEG During Early Adolescence. Sleep, 33(6), 801–809. https://doi.org/10.1093/sleep/33.6.801

Tarokh, L., Van Reen, E., LeBourgeois, M., Seifer, R., & Carskadon, M. A. (2011). Sleep EEG Provides Evidence that Cortical Changes Persist into Late Adolescence. Sleep, 34(10), 1385–1393. https://doi.org/10.5665/SLEEP.1284

Thapar, A., Cooper, M., & Rutter, M. (2017). Neurodevelopmental disorders. The Lancet. Psychiatry, 4(4), 339–346. https://doi.org/10.1016/S2215-0366(16)30376-5

Tononi, G., & Cirelli, C. (2014). Sleep and the Price of Plasticity: From Synaptic and Cellular Homeostasis to Memory Consolidation and Integration. Neuron, 81(1), 12–34. https://doi.org/10.1016/j.neuron.2013.12.025

Vandenbosch, M. M. L. J. Z., van ‘t Ent, D., Boomsma, D. I., Anokhin, A. P., & Smit, D. J. A. (2019). EEG-based age-prediction models as stable and heritable indicators of brain maturational level in children and adolescents. Human Brain Mapping, 40(6), 1919–1926. https://doi.org/10.1002/hbm.24501

Wang, R., Bakker, J. P., Chervin, R. D., Garetz, S. L., Hassan, F., Ishman, S. L., Mitchell, R. B., Morrical, M. G., Naqvi, S. K., Radcliffe, J., Riggan, E. I., Rosen, C. L., Ross, K., Rueschman, M., Tapia, I. E., Taylor, H. G., Zopf, D. A., & Redline, S. (2020). Pediatric Adenotonsillectomy Trial for Snoring (PATS): Protocol for a randomised controlled trial to evaluate the effect of adenotonsillectomy in treating mild obstructive sleep-disordered breathing. BMJ Open, 10(3), e033889. https://doi.org/10.1136/bmjopen-2019-033889

Weinstock, T. G., Rosen, C. L., Marcus, C. L., Garetz, S., Mitchell, R. B., Amin, R., Paruthi, S., Katz, E., Arens, R., Weng, J., Ross, K., Chervin, R. D., Ellenberg, S., Wang, R., & Redline, S. (2014). Predictors of Obstructive Sleep Apnea Severity in Adenotonsillectomy Candidates. Sleep, 37(2), 261–269. https://doi.org/10.5665/sleep.3394

Whitford, T. J., Rennie, C. J., Grieve, S. M., Clark, C. R., Gordon, E., & Williams, L. M. (2006). Brain maturation in adolescence: Concurrent changes in neuroanatomy and neurophysiology. Human Brain Mapping, 28(3), 228–237. https://doi.org/10.1002/hbm.20273

Xu, K., Li, S., Muskens, I. S., Elliott, N., Myint, S. S., Pandey, P., Hansen, H. M., Morimoto, L. M., Kang, A. Y., Ma, X., Metayer, C., Mueller, B. A., Roberts, I., Walsh, K. M., Horvath, S., Wiemels, J. L., & de Smith, A. J. (2022). Accelerated epigenetic aging in newborns with Down syndrome. Aging Cell, 21(7), e13652. https://doi.org/10.1111/acel.13652

