## Supplementary material for "Mapping Typical and Altered Neurodevelopment with Sleep Macro- and Micro-Architecture": Supp. Material

### Supplementary table 1

| Disorders | N | Diagnosis name | Diagnosis ICD codes |
|---|---|---|---|
| ASD | 196 | Autistic disorder; Autistic disorder, current or active state | F84.0; 299.00 |
| ADHD | 525 | Attention-deficit hyperactivity disorder, combined type; Attention deficit disorder with hyperactivity(314.01); Attention-deficit hyperactivity disorder, unspecified type; Attention-deficit hyperactivity disorder, predominantly inattentive type; Attention-deficit hyperactivity disorder, other type; Attention deficit disorder without mention of hyperactivity; Attention-deficit hyperactivity disorder, predominantly hyperactive type | F90.2; 314.01; F90.9; F90.0; F90.8; 314.00; F90.1 |
| IntDis | 167 | Unspecified intellectual disabilities; Moderate intellectual disabilities; Mild intellectual disabilities; Severe intellectual disabilities; Profound intellectual disabilities; Other intellectual disabilities | F79; F71; F70; F72; 319; 318.2; 317; F78; 318.1; F73; 318.0 |
| DS | 140 | Down syndrome, unspecified; Down's syndrome; Trisomy 21, mosaicism (mitotic nondisjunction); Trisomy 21, translocation; Trisomy 21, nonmosaicism (meiotic nondisjunction) | Q90.9; 758.0; Q90.1; Q90.2; Q90.0 |
| CP | 138 | Spastic quadriplegic cerebral palsy; Cerebral palsy, unspecified; Other cerebral palsy; Spastic diplegic cerebral palsy; Spastic hemiplegic cerebral palsy; Infantile cerebral palsy, unspecified; Other specified infantile cerebral palsy; Athetoid cerebral palsy; Ataxic cerebral palsy | G80.0; G80.9; G80.8; G80.1; G80.2; 343.9; 343.8; G80.3; 333.71; G80.4 |
| Epilepsy | 242 | Epilepsy, unspecified, not intractable, without status epilepticus; Localization-related (focal) (partial) symptomatic epilepsy and epileptic syndromes with complex partial seizures, not intractable, with status epilepticus; Generalized idiopathic epilepsy and epileptic syndromes, not intractable, without status epilepticus; Localization-related (focal) (partial) symptomatic epilepsy and epileptic syndromes with complex partial seizures, not intractable, without status epilepticus; Epileptic spasms, not intractable, without status epilepticus; Localization-related (focal) (partial) symptomatic epilepsy and epileptic syndromes with simple partial seizures, not intractable, without status epilepticus; Todd's paralysis (postepileptic); Lennox-Gastaut syndrome, intractable, with status epilepticus; Localization-related (focal) (partial) symptomatic epilepsy and epileptic syndromes with complex partial seizures, intractable, without status epilepticus; Generalized idiopathic epilepsy and epileptic syndromes, intractable, without status epilepticus; Lennox-Gastaut syndrome, intractable, without status epilepticus; Absence epileptic syndrome, not intractable, without status epilepticus; Other generalized epilepsy and epileptic syndromes, not intractable, without status epilepticus; Absence epileptic syndrome, intractable, without status epilepticus; Other generalized epilepsy and epileptic syndromes, intractable, without status epilepticus; Localization-related (focal) (partial) symptomatic epilepsy and epileptic syndromes with simple partial seizures, intractable, without status epilepticus; Epilepsy, unspecified, intractable, without status epilepticus; Other epilepsy, intractable, without status epilepticus; Lennox-Gastaut syndrome, not intractable, without status epilepticus; Epileptic spasms, intractable, with status epilepticus; Localization-related (focal) (partial) symptomatic epilepsy and epileptic syndromes with complex partial seizures, intractable, with status epilepticus; Localization-related (focal) (partial) idiopathic epilepsy and epileptic syndromes with seizures of localized onset, intractable, without status epilepticus; Epilepsy, unspecified, not intractable, with status epilepticus; Other epilepsy, not intractable, without status epilepticus; Epileptic spasms, intractable, without status epilepticus; Unspecified epilepsy without mention of intractable epilepsy; Epilepsy, unspecified, intractable, with status epilepticus; Localization-related (focal) (partial) epilepsy and epileptic syndromes with complex partial seizures, without mention of intractable epilepsy; Infantile spasms with intractable epilepsy; Other epilepsy, not intractable, with status epilepticus; Localization-related (focal) (partial) idiopathic epilepsy and epileptic syndromes with seizures of localized onset, not intractable, without status epilepticus; Localization-related (focal) (partial) idiopathic epilepsy and epileptic syndromes with seizures of localized onset, not intractable, with status epilepticus; Infantile spasms without mention of intractable epilepsy; Unspecified epilepsy with intractable epilepsy; Localization-related (focal) (partial) epilepsy and epileptic syndromes with simple partial seizures, without mention of intractable epilepsy; Generalized nonconvulsive epilepsy without mention of intractable epilepsy; Other forms of epilepsy and recurrent seizures without mention of intractable epilepsy; Generalized convulsive epilepsy without mention of intractable epilepsy; Generalized convulsive epilepsy with intractable epilepsy; Localization-related (focal) (partial) epilepsy and epileptic syndromes with complex partial seizures, with intractable epilepsy; Generalized nonconvulsive epilepsy with intractable epilepsy; Epileptic grand mal status; Generalized idiopathic epilepsy and epileptic syndromes, not intractable, with status epilepticus; Other generalized epilepsy and epileptic syndromes, not intractable, with status epilepticus; Other generalized epilepsy and epileptic syndromes, intractable, with status epilepticus; Epileptic petit mal status; Localization-related (focal) (partial) symptomatic epilepsy and epileptic syndromes with simple partial seizures, not intractable, with status epilepticus; Epileptic seizures related to external causes, not intractable, with status epilepticus; Other epilepsy, intractable, with status epilepticus; Localization-related (focal) (partial) symptomatic epilepsy and epileptic syndromes with simple partial seizures, intractable, with status epilepticus; Generalized idiopathic epilepsy and epileptic syndromes, intractable, with status epilepticus; Lennox-Gastaut syndrome, not intractable, with status epilepticus; Localization-related (focal) (partial) epilepsy and epileptic syndromes with simple partial seizures, with intractable epilepsy; Juvenile myoclonic epilepsy, intractable, without status epilepticus; Other forms of epilepsy and recurrent seizures with intractable epilepsy; Absence epileptic syndrome, not intractable, with status epilepticus; Epileptic seizures related to external causes, not intractable, without status epilepticus; Localization-related (focal) (partial) idiopathic epilepsy and epileptic syndromes with seizures of localized onset, intractable, with status epilepticus; Epileptic spasms, not intractable, with status epilepticus | G40.909; G40.201; G40.309; G40.209; G40.822; G40.109; G83.84; G40.813; G40.219; G40.319; G40.814; G40.A09; G40.409; G40.A19; G40.419; G40.119; G40.919; G40.804; G40.812; G40.823; G40.211; G40.019; G40.901; G40.802; G40.824; 345.90; G40.911; 345.40; 345.61; G40.801; G40.009; G40.001; 345.60; 345.91; 345.50; 345.00; 345.80; 345.10; 345.11; 345.41; 345.01; 345.3; G40.301; G40.401; G40.411; 345.2; G40.101; G40.501; G40.803; G40.111; G40.311; G40.811; 345.51; G40.B19; 345.81; G40.A01; G40.509; G40.011; G40.821 |
| Central apnea | 58 | Primary central sleep apnea; Central sleep apnea in conditions classified elsewhere(327.27); Central sleep apnea in conditions classified elsewhere | G47.31; 327.21; 327.27; G47.37 |
| Other apneas | 2008 | Obstructive sleep apnea (adult) (pediatric); Sleep apnea, unspecified; Apnea, not elsewhere classified; Other sleep apnea; Unspecified sleep apnea; Apnea; Primary apnea of newborn; Other apnea of newborn; Other organic sleep apnea; Organic sleep apnea, unspecified | G47.33; G47.30; R06.81; G47.39; 327.23; 780.57; 786.03; 770.81; P28.4; 770.82; 327.29; 327.20 |
| Insomnia | 403 | Insomnia due to medical condition; Insomnia due to other mental disorder; Insomnia, unspecified; Other insomnia; Problems related to behavioral insomnia of childhood; Psychophysiologic insomnia; Behavioral insomnia of childhood, sleep-onset association type; Behavioral insomnia of childhood, unspecified type; Behavioral insomnia of childhood, combined type; Primary insomnia; Behavioral insomnia of childhood, limit setting type; Insomnia due to medical condition classified elsewhere; Other insomnia not due to a substance or known physiological condition; Adjustment insomnia | G47.01; F51.05; G47.00; G47.09; 780.52; V69.5; F51.04; Z73.810; Z73.819; Z73.812; F51.01; Z73.811; 327.01; F51.09; F51.02 |
| Hypersomnia | 175 | Hypersomnia, unspecified; Other hypersomnia; Idiopathic hypersomnia with long sleep time; Hypersomnia due to medical condition; Idiopathic hypersomnia without long sleep time | G47.10; G47.19; G47.11; 780.54; G47.14; G47.12 |
| Other sleep disorders | 1650 | Sleep disorder, unspecified; Sleep terrors (night terrors); Restless legs syndrome; Sleep deprivation; Other sleep disorders; Other sleep disturbances; Sleep disturbance, unspecified; Restless legs syndrome (RLS); Circadian rhythm sleep disorder, unspecified type; Parasomnia, unspecified; Sleep related hypoventilation/hypoxemia in conditions classifiable elsewhere; Problems related to lack of adequate sleep; Sleep related bruxism; Sleepwalking (somnambulism); Idiopathic sleep related nonobstructive alveolar hypoventilation; Dysfunctions associated with sleep stages or arousal from sleep; Other dysfunctions of sleep stages or arousal from sleep; Sleep arousal disorder; Inadequate sleep hygiene; Circadian rhythm sleep disorder, irregular sleep wake type; | G47.9; G25.81; 333.94; Z72.820; G47.8; F51.8; 780.59; 780.50; G47.20; G47.50; 327.26; V69.4; G47.63; 327.53; G47.34; 327.24; 780.56; 307.47; 307.46; Z72.821; |

| | |
|---|---|
| Other psychoactive substance use, unspecified with psychoactive substance-induced sleep disorder; Other parasomnia; Persistent disorder of initiating or maintaining sleep; Sleep related hypoventilation in conditions classified elsewhere; Other sleep disorders not due to a substance or known physiological condition; Other specific disorder of sleep of nonorganic origin; Sleep disorder not due to a substance or known physiological condition, unspecified; Insufficient sleep syndrome; Circadian rhythm sleep disorder, delayed sleep phase type; Other sleep related movement disorders; Circadian rhythm sleep disorder of nonorganic origin; Circadian rhythm sleep disorder, unspecified; Sleep related movement disorder, unspecified; Recurrent isolated sleep paralysis; Circadian rhythm sleep disorder, free running type; REM sleep behavior disorder; Parasomnia in conditions classified elsewhere; Other organic sleep disorders; Nonorganic sleep disorder, unspecified; Sleep related leg cramps | G47.23; F19.982; G47.59; 307.42; G47.36; 307.49; F51.9; F51.12; G47.21; G47.69; 307.45; 327.30; 780.58; G47.53; G47.24; G47.52; 327.42; G47.54; 327.8; 307.40; G47.62 |

### Supplementary table 2 Medication overlapping PSG recording in NCH sample

| Therapeutical class | Pharmaceutic class | Therapeutic subclass | ASD | ADHD | IntDis | DS | CP | Epilepsy | Rest |
|---|---|---|---|---|---|---|---|---|---|
| ANTIHISTAMINES | | | 27 (14%) | 76 (14%) | 34 (20%) | 12 (9%) | 26 (19%) | 43 (18%) | 157 (9%) |
| | ANTIHISTAMINES - 2ND GENERATION | cetirizine HCl; loratadine; fexofenadine HCl | 15 (8%) | 46 (9%) | 20 (12%) | 9 (6%) | 15 (11%) | 24 (10%) | 116 (6%) |
| | ANTIHISTAMINES - 1ST GENERATION | hydroxyzine HCl; cyproheptadine HCl; di phenhydramine HCl; ; hydroxyzine pamo ate; promethazine HCl | 14 (7%) | 35 (7%) | 20 (12%) | 5 (4%) | 14 (10%) | 23 (10%) | 50 (3%) |
| | EYE ANTIHISTAMINES | azelastine HCl; ketotifen fumarate | 1 (1%) | 2 (0%) | 1 (1%) | 0 (0%) | 2 (1%) | 1 (0%) | 12 (1%) |
| PSYCHOTHERAPEUTIC DRUGS | | | 52 (27%) | 160 (30%) | 51 (31%) | 8 (6%) | 19 (14%) | 41 (17%) | 51 (3%) |
| | SELECTIVE SEROTONIN REUPTAKE INHIBITOR (SSRIS) | citalopram hydrobromide; fluoxetine HCl ; escitalopram oxalate; sertraline HCl; flu voxamine maleate | 24 (12%) | 73 (14%) | 19 (11%) | 3 (2%) | 2 (1%) | 12 (5%) | 32 (2%) |
| | ALPHA-2 RECEPTOR ANTAGONIST ANTIDEPRESSANTS | mirtazapine | 0 (0%) | 4 (1%) | 0 (0%) | 0 (0%) | 0 (0%) | 1 (0%) | 2 (0%) |
| | ANTIPSYCHOTIC,ATYPICAL,DOPAMINE,SEROTONIN A NTAGNST | risperidone; quetiapine fumarate; lurasi done HCl; olanzapine | 13 (7%) | 26 (5%) | 10 (6%) | 2 (1%) | 1 (1%) | 6 (2%) | 3 (0%) |
| | TRICYCLIC ANTIDEPRESSANTS,REL.NON-SEL.REUPT-INHIB | amitriptyline HCl; doxepin HCl; ; clomipr amine HCl | 1 (1%) | 3 (1%) | 0 (0%) | 0 (0%) | 0 (0%) | 2 (1%) | 11 (1%) |
| | ANTIPSYCHOTICS, ATYP, D2 PARTIAL AGONIST/5HT M IXED | aripiprazole | 4 (2%) | 14 (3%) | 4 (2%) | 1 (1%) | 0 (0%) | 2 (1%) | 2 (0%) |
| | TX FOR ADHD - SELECTIVE ALPHA-2 RECEPTOR AGONIST | guanfacine HCl; clonidine HCl | 9 (5%) | 35 (7%) | 6 (4%) | 1 (1%) | 0 (0%) | 1 (0%) | 4 (0%) |
| | SEROTONIN-2 ANTAGONIST/REUPTAKE INHIBITORS (SARIS) | trazodone HCl | 2 (1%) | 14 (3%) | 5 (3%) | 0 (0%) | 3 (2%) | 6 (2%) | 2 (0%) |
| | NARCOLEPSY AND SLEEP DISORDER THERAPY AGENTS | modafinil; armodafinil | 12 (6%) | 59 (11%) | 11 (7%) | 2 (1%) | 2 (1%) | 4 (2%) | 1 (0%) |
| | TX FOR ATTENTION DEFICIT-HYPERACT(ADHD)/NARCOLEPSY | methylphenidate HCl; dexmethylphenid ate HCl | 8 (4%) | 2 (0%) | 13 (8%) | 1 (1%) | 14 (10%) | 18 (7%) | 2 (0%) |
| | ANTI-ANXIETY - BENZODIAZEPINES | diazepam; lorazepam | 4 (2%) | 11 (2%) | 2 (1%) | 0 (0%) | 0 (0%) | 1 (0%) | 0 (0%) |
| | TX FOR ATTENTION DEFICIT-HYPERACT.(ADHD), NRI-TYPE | atomoxetine HCl | 5 (3%) | 16 (3%) | 4 (2%) | 2 (1%) | 0 (0%) | 3 (1%) | 0 (0%) |
| | ADRENERGICS, AROMATIC, NON-CATECHOLAMINE | lisdexamfetamine dimesylate | 0 (0%) | 1 (0%) | 0 (0%) | 0 (0%) | 0 (0%) | 0 (0%) | 3 (0%) |
| | ANTI-ANXIETY DRUGS | buspirone HCl | 2 (1%) | 1 (0%) | 1 (1%) | 0 (0%) | 0 (0%) | 0 (0%) | 0 (0%) |
| | BIPOLAR DISORDER DRUGS | lithium carbonate | 2 (1%) | 2 (0%) | 2 (1%) | 0 (0%) | 0 (0%) | 0 (0%) | 4 (0%) |
| | SEROTONIN-NOREPINEPHRINE REUPTAKE-INHIB (SNRIS) | venlafaxine HCl; duloxetine HCl | 1 (1%) | 1 (0%) | 0 (0%) | 0 (0%) | 0 (0%) | 0 (0%) | 0 (0%) |
| | NOREPINEPHRINE AND DOPAMINE REUPTAKE INHIB ( NDRIS) | bupropion HCl | 32 (16%) | 40 (8%) | 52 (31%) | 11 (8%) | 58 (42%) | 132 (55%) | 17 (1%) |
| CNS DRUGS | | | 14 (7%) | 7 (1%) | 23 (14%) | 6 (4%) | 32 (23%) | 60 (25%) | 2 (0%) |
| | ANTICONVULSANT - BENZODIAZEPINE TYPE | clobazam; diazepam; clonazepam | 25 (13%) | 34 (6%) | 44 (26%) | 10 (7%) | 50 (36%) | 115 (48%) | 16 (1%) |
| | ANTICONVULSANTS | divalproex sodium; zonisamide; oxcarba zepine; valproic acid (as sodium salt); ; t opiramate; rufinamide; lacosamide; preg abalin; gabapentin; lamotrigine; brivarac etam; ethosuximide; levetiracetam; felb | 4 (2%) | 1 (0%) | 3 (2%) | 0 (0%) | 2 (1%) | 6 (2%) | 0 (0%) |

| | | | | | | | | | |
|---|---|---|---|---|---|---|---|---|---|
| | | amate; vigabatrin; carbamazepine; phenytoin; perampanel; diazepam | | | | | | | |
| | ANTICONVULSANT - CANNABINOID TYPE | cannabidiol (CBD) | 25 (13%) | 62 (12%) | 28 (17%) | 5 (4%) | 16 (12%) | 31 (13%) | 73 (4%) |
| HORMONES | | | 3 (2%) | 3 (1%) | 4 (2%) | 1 (1%) | 3 (2%) | 7 (3%) | 8 (0%) |
| | GROWTH HORMONES | somatropin | 17 (9%) | 46 (9%) | 19 (11%) | 4 (3%) | 10 (7%) | 21 (9%) | 18 (1%) |
| | PINEAL HORMONE AGENTS | melatonin; | 3 (2%) | 9 (2%) | 3 (2%) | 2 (1%) | 3 (2%) | 8 (3%) | 39 (2%) |
| | GLUCOCORTICOIDS | prednisone; hydrocortisone; hydrocortisone sodium succ/PF; prednisolone sodium phosphate; hydrocortisone sod succinate; deflazacort; methylprednisolone; dexamethasone sodium phosphate | 1 (1%) | 5 (1%) | 3 (2%) | 0 (0%) | 0 (0%) | 1 (0%) | 2 (0%) |
| | ANTIDIURETIC AND VASOPRESSOR HORMONES | desmopressin acetate | 1 (1%) | 3 (1%) | 1 (1%) | 0 (0%) | 1 (1%) | 2 (1%) | 5 (0%) |
| | PROGESTATIONAL AGENTS | norethindrone acetate; medroxyprogesterone acetate | 1 (1%) | 1 (0%) | 1 (1%) | 0 (0%) | 1 (1%) | 1 (0%) | 0 (0%) |
| | LHRH(GNRH)AGNST PIT.SUP-CENTRAL PRECOCIOUS PUBERTY | leuprolide acetate | 0 (0%) | 0 (0%) | 0 (0%) | 0 (0%) | 0 (0%) | 0 (0%) | 5 (0%) |
| | MINERALOCORTICOIDS | fludrocortisone acetate | 0 (0%) | 0 (0%) | 1 (1%) | 0 (0%) | 0 (0%) | 0 (0%) | 0 (0%) |
| | ESTROGENIC AGENTS | estradiol | 1 (1%) | 0 (0%) | 2 (1%) | 0 (0%) | 2 (1%) | 4 (2%) | 1 (0%) |
| SEDATIVE/HYPNOTICS | | | 1 (1%) | 0 (0%) | 1 (1%) | 0 (0%) | 2 (1%) | 3 (1%) | 0 (0%) |
| | BARBITURATES | phenobarbital | 0 (0%) | 0 (0%) | 1 (1%) | 0 (0%) | 0 (0%) | 1 (0%) | 1 (0%) |
| | SEDATIVE-HYPNOTICS - BENZODIAZEPINES | lorazepam | 0 (0%) | 0 (0%) | 1 (1%) | 0 (0%) | 0 (0%) | 1 (0%) | 1 (0%) |

Supplementary table 3 Differences in absolute power between subgroups and the rest of the NCH sample

| STAGE & BAND | | ASD | | | ADHD | | | Intellectual disabilities | | | Down Syndrome | | | Cerebral palsy | | | Epilepsy | | |
|---|---|---|---|---|---|---|---|---|---|---|---|---|---|---|---|---|---|---|---|
| | | Max b | Min p | CHS, p < 0.05 | Max b | Min p | CHS, p < 0.05 | Max b | Min p | CHS, p < 0.05 | Max b | Min p | CHS, p < 0.05 | Max b | Min p | CHS, p < 0.05 | Max b | Min p | CHS, p < 0.05 |
| N2 | SLOW | 0.17 | 0.04 | F4 | - | - | - | -0.21 | 0.03 | F4 | 0.45 | 7E-09 | all | - | - | - | 0.19 | 0.01 | O2 |
| | DELTA | 0.24 | 0.002 | F3, F4 | - | - | - | -0.27 | 0.005 | F4 | 0.35 | 5E-07 | O1, O2 | - | - | - | 0.18 | 0.009 | O2 |
| | THETA | 0.14 | 0.02 | F4 | - | - | - | -0.19 | 0.01 | F4 | 0.35 | 9E-07 | all | - | - | - | 0.14 | 0.04 | O2 |
| | ALPHA | - | - | - | - | - | - | - | - | - | 0.31 | 0.001 | F3, F4, O1, O2 | - | - | - | 0.22 | 0.02 | F3, O2 |
| | SIGMA | - | - | - | 0.16 | 0.003 | F3, F4, C3, C4 | - | - | - | 0.28 | 0.003 | O1, O2 | - | - | - | 0.33 | 4E-04 | all |
| | BETA | - | - | - | - | - | - | - | - | - | 0.94 | 2E-24 | all | 0.29 | 0.008 | C3 | 0.29 | 0.002 | F3, F4, C3, C4 |
| | TOTAL | 0.19 | 0.01 | F3, F4 | - | - | - | -0.27 | 0.005 | F4 | 0.4 | 3E-08 | all, but C4 | - | - | - | 0.2 | 0.004 | O2 |
| N3 | SLOW | - | - | - | - | - | - | -0.2 | 0.02 | O2 | 0.98 | 3E-26 | all | - | - | - | - | - | - |
| | DELTA | - | - | - | - | - | - | -0.2 | 0.02 | F4, C4, O2 | 0.59 | 1E-16 | all | - | - | - | - | - | - |
| | THETA | 0.14 | 0.03 | F4 | - | - | - | - | - | - | 0.39 | 2E-08 | all | - | - | - | - | - | - |
| | ALPHA | - | - | - | - | - | - | - | - | - | 0.44 | 9E-07 | C3, C4, O1, O2 | - | - | - | 0.18 | 0.05 | F3 |
| | SIGMA | - | - | - | 0.15 | 0.01 | F3, F4, C4 | - | - | - | 0.3 | 0.002 | O1, O2 | - | - | - | 0.29 | 0.003 | all |
| | BETA | - | - | - | - | - | - | 0.22 | 0.03 | F3 | 0.43 | 8E-06 | all, but C4 | - | - | - | 0.26 | 0.004 | F3, F4 |
| | TOTAL | 0.16 | 0.05 | F4 | - | - | - | -0.18 | 0.03 | C4, O2 | 0.63 | 2E-18 | all | - | - | - | - | - | - |
| R | SLOW | - | - | - | -0.09 | 0.05 | O2 | - | - | - | 0.82 | 3E-26 | all | - | - | - | 0.19 | 0.009 | all, but O1 |
| | DELTA | 0.11 | 0.03 | F4 | - | - | - | 0.15 | 0.03 | O2 | 0.96 | 1E-55 | all | 0.16 | 0.02 | C3 | 0.21 | 3E-04 | all, but O1 |
| | THETA | - | - | - | - | - | - | - | - | - | 0.77 | 2E-27 | all | - | - | - | 0.14 | 0.04 | C4 |
| | ALPHA | - | - | - | - | - | - | - | - | - | 0.61 | 2E-11 | all, but O2 | -0.23 | 0.05 | O2 | - | - | - |
| | SIGMA | 0.2 | 0.02 | F3, F4, C3 | - | - | - | - | - | - | 0.79 | 5E-17 | all | - | - | - | 0.3 | 0.002 | F3, F4, C4 |
| | BETA | 0.18 | 0.03 | F3 | - | - | - | - | - | - | 0.94 | 5E-23 | all | - | - | - | 0.33 | 6E-04 | F3, F4, C3, C4 |
| | TOTAL | 0.12 | 0.02 | F4 | - | - | - | - | - | - | 1 | 1E-53 | all | 0.15 | 0.04 | C4 | 0.21 | 5E-04 | all, but O1 |

Supplementary table 4 Differences in relative power between subgroups and the rest of the NCH sample

| STAGE & BAND | | ASD | | | ADHD | | | Intellectual disabilities | | | Down Syndrome | | | Cerebral palsy | | | Epilepsy | | |
|---|---|---|---|---|---|---|---|---|---|---|---|---|---|---|---|---|---|---|---|
| | | Max b | Min p | CHS, p < 0.05 | Max b | Min p | CHS, p < 0.05 | Max b | Min p | CHS, p < 0.05 | Max b | Min p | CHS, p < 0.05 | Max b | Min p | CHS, p < 0.05 | Max b | Min p | CHS, p < 0.05 |
| N2 | SLOW | - | - | - | 0.14 | 0.01 | F3, C3 | -0.23 | 0.03 | C4 | 0.38 | 1E-04 | O2 | -0.35 | 0.002 | F3, C3, O1 | - | - | - |
| | DELTA | - | - | - | - | - | - | - | - | - | -0.6 | 2E-09 | all, but O2 | 0.24 | 0.03 | O1 | - | - | - |
| | THETA | 0.2 | 0.02 | O1 | -0.12 | 0.006 | F3, F4, C3 | - | - | - | 0.31 | 5E-04 | F3, F4, C3, C4 | - | - | - | - | - | - |
| | ALPHA | - | - | - | 0.13 | 0.03 | F4 | - | - | - | -0.59 | 1E-09 | all | - | - | - | - | - | - |
| | SIGMA | - | - | - | - | - | - | - | - | - | - | - | - | - | - | - | - | - | - |
| | BETA | - | - | - | - | - | - | 0.21 | 0.04 | F4, C4 | 0.55 | 6E-08 | all, but O2 | - | - | - | - | - | - |
| N3 | SLOW | - | - | - | - | - | - | 0.22 | 0.03 | F3 | 0.94 | 1E-21 | all | - | - | - | - | - | - |
| | DELTA | - | - | - | 0.14 | 0.01 | F4, C3 | -0.21 | 0.03 | O1 | -0.68 | 9E-13 | F3, F4, C3, C4 | - | - | - | -0.22 | 0.02 | F4, C4, O2 |
| | THETA | 0.17 | 0.04 | O2 | -0.1 | 0.04 | F3 | - | - | - | -0.48 | 7E-07 | C4, O1, O2 | - | - | - | - | - | - |
| | ALPHA | - | - | - | - | - | - | - | - | - | -0.51 | 3E-07 | all | - | - | - | 0.24 | 0.01 | F3, C3, C4 |
| | SIGMA | - | - | - | - | - | - | - | - | - | -0.45 | 1E-08 | all | -0.25 | 0.03 | F3, F4 | 0.28 | 2E-04 | all |
| | BETA | - | - | - | -0.11 | 0.05 | C3 | 0.36 | 3E-04 | all, but O1 | -0.39 | 3E-05 | all | - | - | - | 0.26 | 0.002 | all |
| R | SLOW | - | - | - | - | - | - | - | - | - | 0.71 | 8E-13 | C4, O1, O2 | - | - | - | - | - | - |
| | DELTA | -0.16 | 0.01 | O2 | - | - | - | 0.19 | 0.02 | C3, O2 | 0.58 | 3E-14 | all | - | - | - | - | - | - |
| | THETA | - | - | - | - | - | - | -0.2 | 0.05 | O2 | -0.57 | 2E-09 | all | - | - | - | - | - | - |
| | ALPHA | - | - | - | 0.1 | 0.02 | O1 | -0.24 | 0.001 | O1, O2 | -0.64 | 3E-17 | all | - | - | - | - | - | - |
| | SIGMA | 0.19 | 0.001 | C3, O1, O2 | - | - | - | 0.17 | 0.03 | C3 | -0.4 | 8E-08 | all, but O1 | - | - | - | - | - | - |
| | BETA | - | - | - | - | - | - | - | - | - | -0.24 | 0.004 | F3, F4, C3, C4 | - | - | - | - | - | - |

Supplementary table 5 Differences in sleep microarchitecture between subgroups and the rest of the NCH sample

| Metric | | ASD | | | ADHD | | | Intellectual disabilities | | | Down Syndrome | | | Cerebral palsy | | | Epilepsy | | |
|---|---|---|---|---|---|---|---|---|---|---|---|---|---|---|---|---|---|---|---|
| | | Max b | Min p | CHS, $p < 0.05$ | Max b | Min p | CHS, $p < 0.05$ | Max b | Min p | CHS, $p < 0.05$ | Max b | Min p | CHS, $p < 0.05$ | Max b | Min p | CHS, $p < 0.05$ | Max b | Min p | CHS, $p < 0.05$ |
| Density | SS | - | - | - | 0.22 | 3E-05 | F3, F4, C3, C4 | -0.29 | 0.003 | F3, F4 | -0.85 | 2E-20 | all, but O1 | -0.4 | 2E-04 | F3, F4 | - | - | - |
| | FS | - | - | - | - | - | - | -0.21 | 0.02 | F3, O2 | -0.66 | 5E-15 | all | -0.2 | 0.05 | F3 | -0.19 | 0.03 | C3, C4 |
| Amplitude | SS | - | - | - | 0.16 | 0.005 | F3, F4, C3 | - | - | - | 0.36 | 2E-04 | F3, F4, O1, O2 | - | - | - | 0.28 | 0.003 | F3, C4, O2 |
| | FS | - | - | - | - | - | - | - | - | - | 0.43 | 1E-05 | F3, F4, O1, O2 | - | - | - | 0.39 | 5E-05 | all, but C3 |
| Duration | SS | - | - | - | 0.15 | 0.007 | F3, F4 | - | - | - | -1.11 | 1E-31 | all | -0.42 | 1E-04 | all | -0.21 | 0.02 | F4, C3, C4 |
| | FS | - | - | - | - | - | - | - | - | - | -0.7 | 6E-13 | all | -0.27 | 0.01 | O1 | - | - | - |
| Frequency | SS | 0.18 | 0.03 | O2 | 0.16 | 0.004 | F3, C3, C4, O1 | -0.28 | 0.002 | F3, F4 | - | - | - | -0.37 | 5E-04 | all | - | - | - |
| | FS | - | - | - | -0.19 | 2E-04 | all | 0.31 | 7E-04 | all, but O2 | 0.81 | 8E-21 | all | 0.59 | 2E-08 | all | - | - | - |
| Chirp | SS | - | - | - | -0.14 | 0.006 | F3, F4, O1 | 0.37 | 5E-05 | all | 0.86 | 3E-27 | all | 0.31 | 5E-04 | F3, F4, C3, C4 | - | - | - |
| | FS | -0.18 | 0.04 | F3, O2 | - | - | - | - | - | - | 0.89 | 2E-21 | all, but O2 | 0.24 | 0.03 | C3, C4 | - | - | - |
| SO rate | | 0.22 | 0.001 | F3, F4 | - | - | - | - | - | - | -0.75 | 9E-15 | all | 0.21 | 0.03 | C3 | - | - | - |
| SO duration | | -0.2 | 0.03 | C3, O1 | 0.16 | 0.01 | all, but O2 | - | - | - | 0.64 | 5E-07 | F3, O1, O2 | -0.45 | 3E-04 | all, but F4 | - | - | - |
| SO slope | | 0.18 | 0.03 | F4 | - | - | - | -0.21 | 0.03 | F4 | - | - | - | 0.2 | 0.04 | C3 | - | - | - |
| SO amplitude | | - | - | - | - | - | - | - | - | - | - | - | - | - | - | - | - | - | - |
| SO P2P | | - | - | - | - | - | - | - | - | - | - | - | - | - | - | - | - | - | - |
| Coupling magnitude | SS | -0.17 | 0.03 | F3 | 0.15 | 0.004 | F3 | - | - | - | 0.38 | 9E-04 | F3, F4, C3, O1 | - | - | - | - | - | - |
| | FS | - | - | - | -0.12 | 0.05 | O1 | - | - | - | -0.71 | 1E-08 | F3, F4, C3, C4 | -0.24 | 0.04 | C4 | -0.28 | 0.006 | F4 |
| Coupling overlap | SS | - | - | - | - | - | - | - | - | - | -0.24 | 0.05 | F3 | - | - | - | -0.24 | 0.02 | C3, C4 |
| | FS | -0.23 | 0.01 | C3 | - | - | - | - | - | - | -0.56 | 1E-05 | F3, F4, C3, C4 | -0.39 | 0.002 | F4, C3, C4, O1 | - | - | - |
| Coupling angle | SS | - | - | - | - | - | - | 0.27 | 0.02 | F3 | -0.39 | 0.002 | F4, C3, O1 | 0.31 | 0.01 | O1 | - | - | - |
| | FS | 0.18 | 0.05 | O2 | - | - | - | -0.3 | 0.01 | C4 | -0.74 | 2E-08 | F3, F4, C3, C4 | -0.29 | 0.02 | C3 | -0.35 | 6E-04 | F4 |

Supplementary table 6 Differences in sleep macroarchitecture between subgroups and the rest of the NCH sample

| Metric | ASD | | ADHD | | Intellectual disabilities | | Down Syndrome | | Cerebral palsy | | Epilepsy | |
|---|---|---|---|---|---|---|---|---|---|---|---|---|
| | **b** | **p-value** | **b** | **p-value** | **b** | **p-value** | **b** | **p-value** | **b** | **p-value** | **b** | **p-value** |
| **TST** | - | - | - | - | - | - | - | - | -0.39 | 7.6E-05 | - | - |
| **SME** | -0.19 | 0.016 | - | - | - | - | -0.39 | 2.2E-05 | - | - | - | - |
| **R latency** | - | - | 0.17 | 0.00158 | - | - | 0.42 | 4.9E-06 | - | - | 0.22 | 1.5E-02 |
| **WASO** | - | - | - | - | - | - | 0.44 | 2.3E-06 | 0.2 | 5E-02 | - | - |
| **N of sleep cycles** | - | - | -0.18 | 0.00048 | - | - | -0.33 | 9.1E-05 | -0.46 | 9.6E-07 | -0.19 | 2.2E-02 |
| **SFI** | - | - | - | - | - | - | 0.52 | 1.2E-08 | - | - | - | - |
| **Transition index NR-R** | - | - | - | - | - | - | - | - | -0.31 | 1.4E-03 | - | - |
| **Duration of N1** | - | - | - | - | - | - | - | - | - | - | -0.19 | 3.6E-02 |
| **Duration of N2** | - | - | - | - | - | - | - | - | -0.21 | 3.4E-02 | - | - |
| **Duration of N3** | - | - | - | - | - | - | -0.24 | 4.6E-03 | - | - | - | - |
| **Duration of R** | - | - | - | - | -0.27 | 0.00174 | -0.26 | 2.3E-03 | -0.56 | 4.6E-09 | -0.3 | 3.2E-04 |
| **Proportion of N1** | - | - | - | - | - | - | 0.21 | 2.4E-02 | - | - | -0.2 | 2.7E-02 |
| **Proportion of N2** | - | - | - | - | - | - | 0.26 | 2.2E-03 | - | - | - | - |
| **Proportion of N3** | - | - | - | - | - | - | - | - | 0.23 | 2.5E-02 | 0.17 | 4.4E-02 |
| **Proportion of R** | - | - | - | - | -0.32 | 0.00019 | -0.26 | 2.9E-03 | -0.53 | 1.9E-08 | -0.35 | 3.4E-05 |

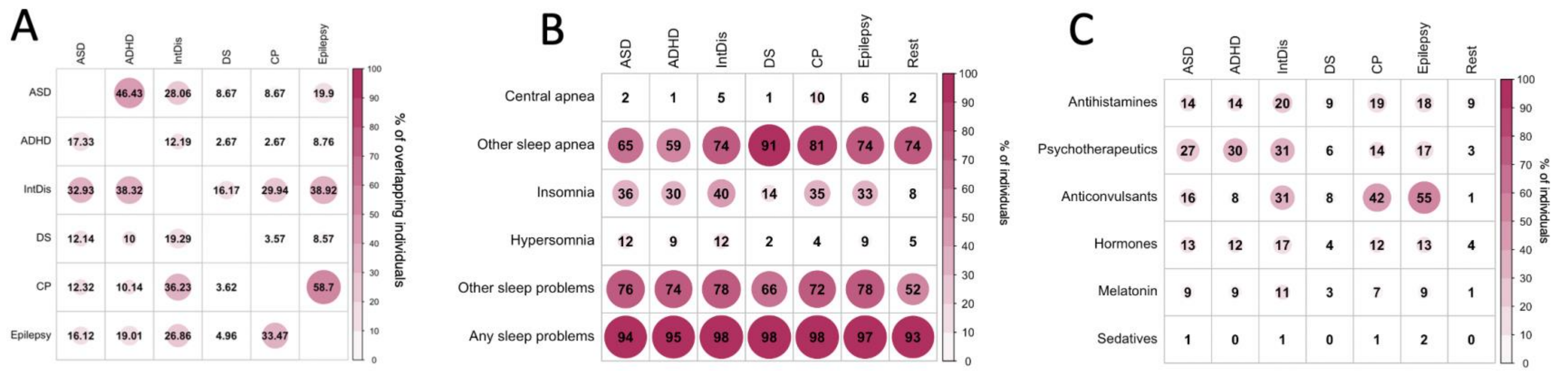


***Figure S1 Sleep-related diagnoses, medication usage in the whole cohort and subgroups and overlap between clinical sub-groups.*** *A – percentage of individuals in a particular NDD subgroup (rows) that had an overlapping NDD diagnosis (columns); B – Percentage of individuals among the NDD subgroups and the rest of the NCH sample (columns of the matrix) that had sleep diagnoses listed as rows in the matrix; C – Percentage of individuals among the NDD subgroups and the rest of the NCH sample (columns of the matrix) who were prescribed medication that could affect sleep (rows in the matrix) over the period overlapping the polysomnography recording.*

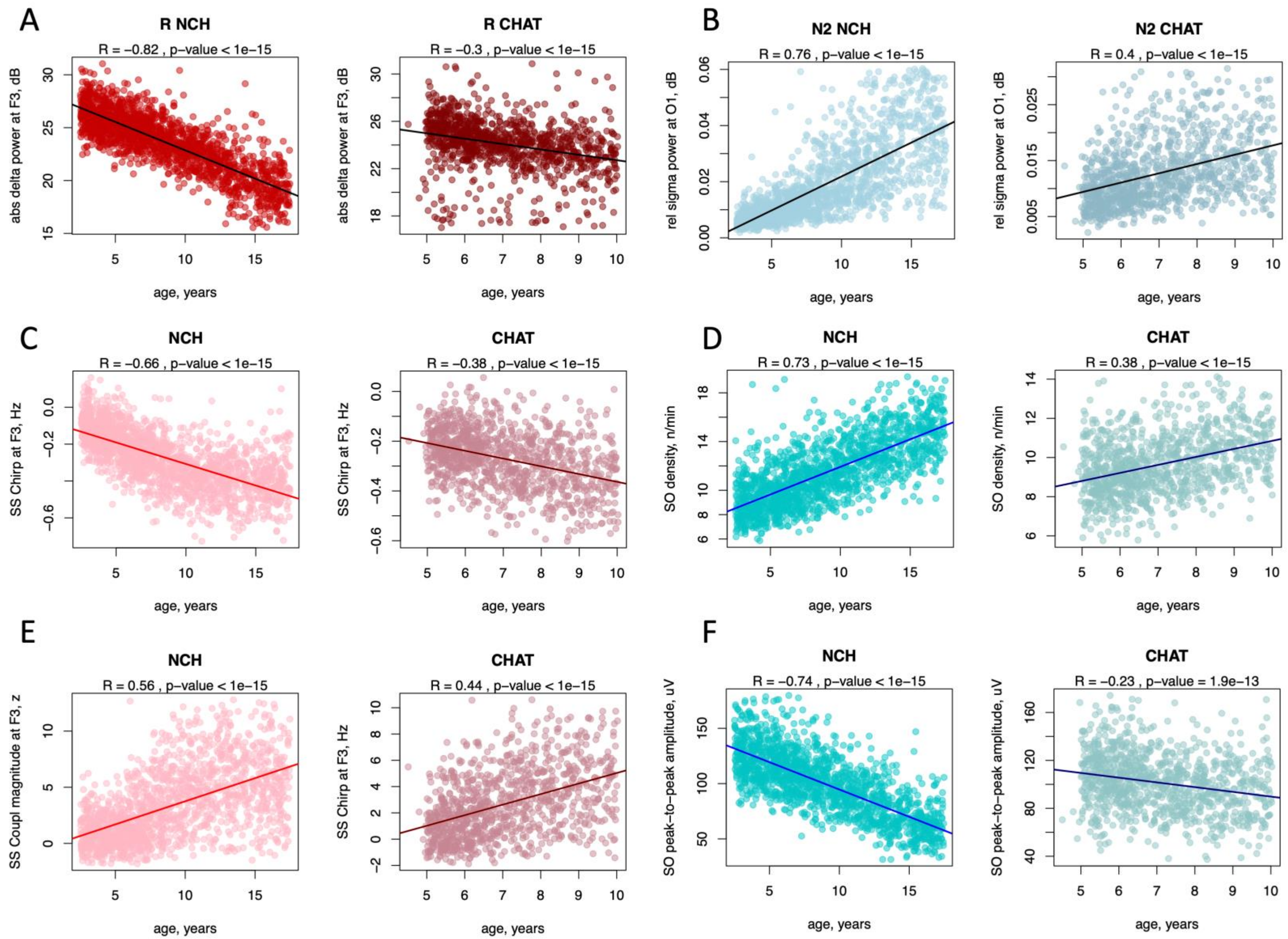


***Figure S2 Sleep micro-architecture estimates with strongest linear age-related changes***

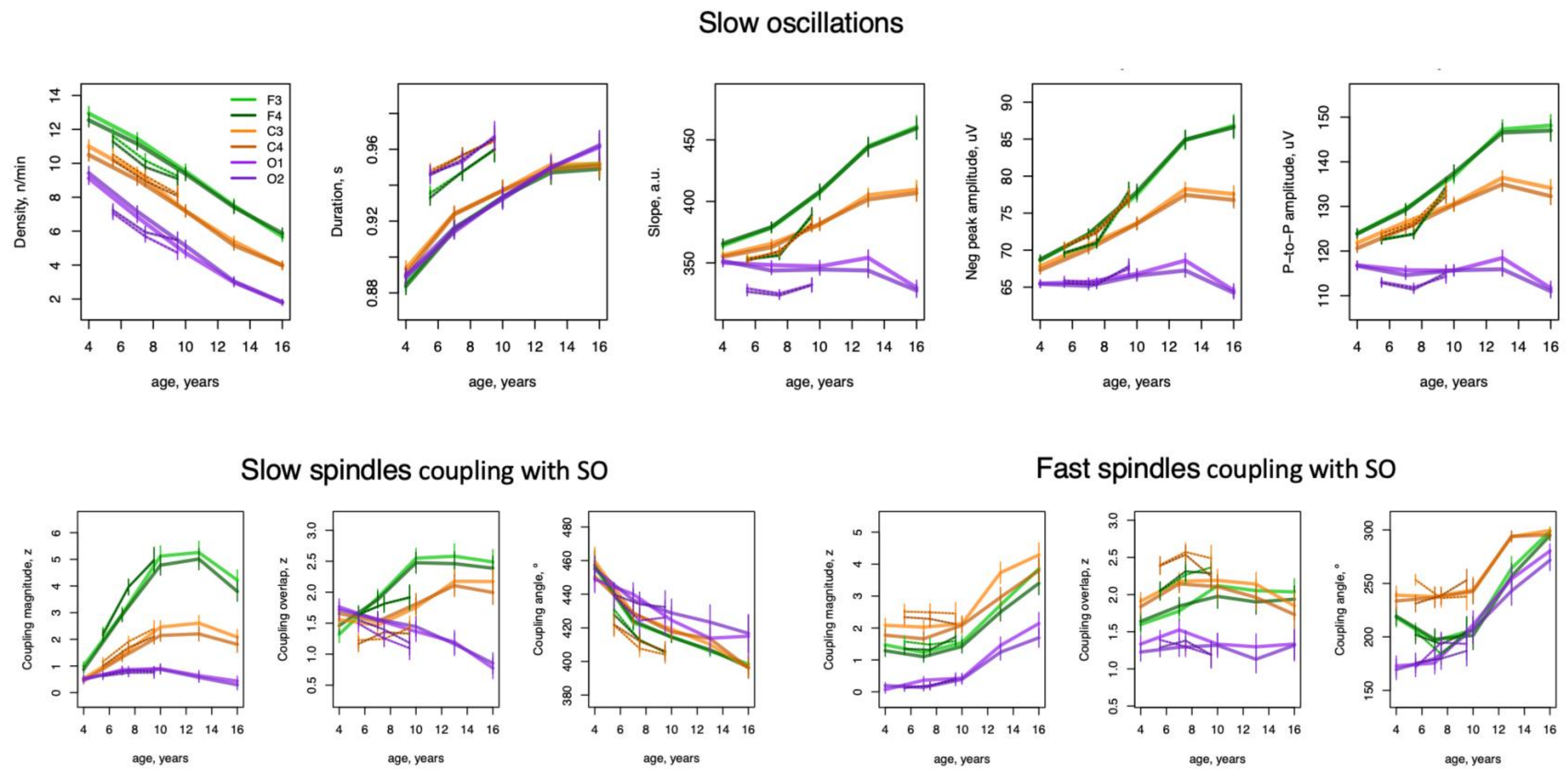


***Sup Figure 3 Age related changes in SO metrics and SO-spindle coupling when SOs detected with absolute amplitude threshold*.**

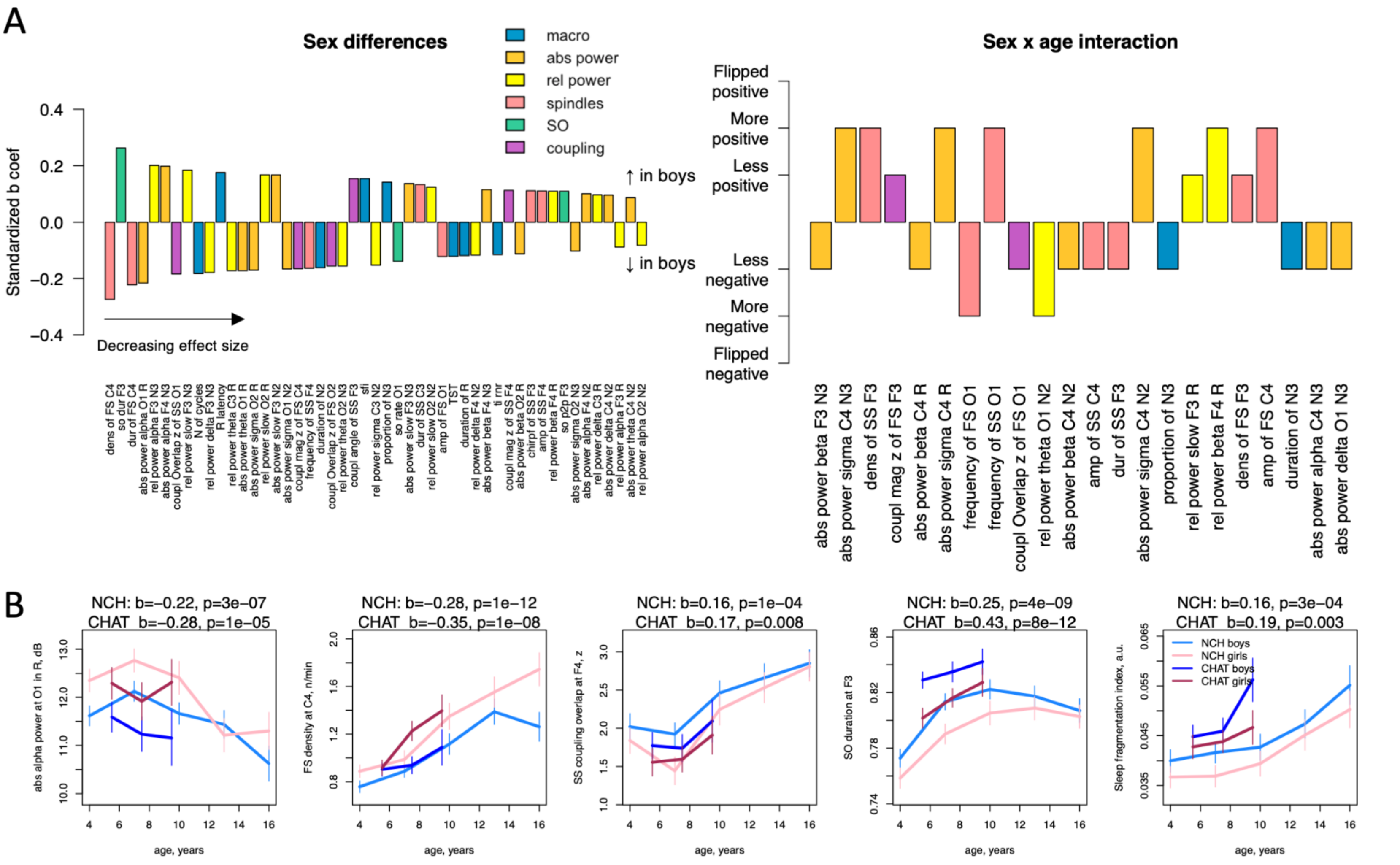


***Figure S4 Sleep parameters with strongest sex effects in NCH sample*** *A – The bar plot on the left represents sorted by the effect size (standardized coefficient from linear regression) significant differences (p <0.05 FDR-corrected) between boys and girls controlling for age and race. Although weaker than age-related effects, we observed sex differences across multiple aspects of sleep architecture. . If similar effects were present across multiple channels, only the one with the largest effect size is plotted. The bar plot on the right illustrates significant sex-by-age effects. To aid interpretation, standardized b-coefficients are transformed to illustrate six possible scenarios: i. male age-trajectory was more positive compared to positive female trajectory (both* $b_m$ *and* $b_f > 0$ *and* $b_m > b_f$*), ii. male age-trajectory was less positive (closer to zero) compared to positive female trajectory (both* $b_m$ *and* $b_f > 0$ *and* $b_m < b_f$*), iii. male age-trajectory was flipped, i.e. significantly (nominal* $p < 0.05$*) positive when female trajectory was negative (*$b_m$ *and* $b_f$ *have different sign and* $b_m > b_f$ *and* $p_m < 0.05$*). B – examples with strongest sex-effects present in both NCH and CHAT samples.*

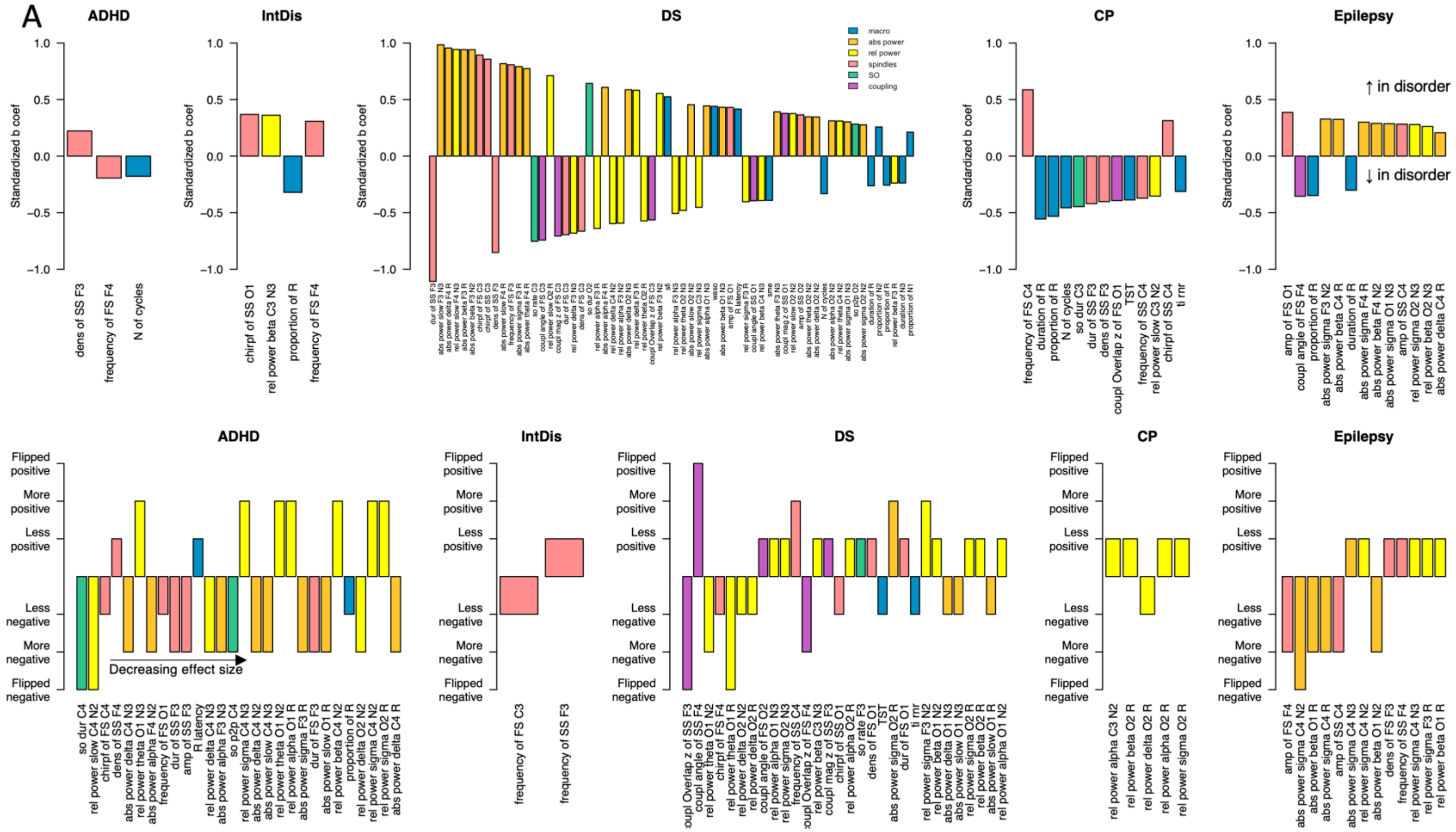


***Figure S5 Alterations in sleep macro and microarchitecture in NDD subgroups*** *The bar plots show significant ($p$<0.05 after FDR correction) subgroup and subgroup-by-age interaction (top and bottom row respectively) ordered by associated absolute standardized b-coefficients from left to right. If similar effects were present across multiple channels, only the one with the largest effect size is plotted. To aid interpretation of subgroup-by-age interaction they are transformed accounting for developmental trajectories in non-NDD sample to illustrate six possible scenarios: i. subgroup age-trajectory was more positive compared to positive non-NDD trajectory (both $b_{subgroup}$ and $b_{non\text{-}NDD} > 0$ and $b_{subgroup} > b_{non\text{-}NDD}$), ii. subgroup age-trajectory was less positive (closer to zero) compared to positive non-NDD trajectory (both $b_{subgroup}$ and $b_{non\text{-}NDD} > 0$ and $b_{subgroup} < b_{non\text{-}NDD}$), iii. subgroup age-trajectory was flipped, i.e. significantly (nominal $p < 0.05$) positive when non-NDD trajectory was negative ($b_{subgroup}$ and $b_{non\text{-}NDD}$ have different sign and $b_{subgroup} > b_{non\text{-}NDD}$ and $p_{subgroup} < 0.05$). Note that ASD group is not plotted because there were no significant affects after multiple comparison correction with FDR method. See* ***Sup.Table 3-5*** *for full list of uncorrected results*
